# SWING regions prime chromatin for nuclear speckle–mediated gene regulation

**DOI:** 10.1101/2025.10.01.679801

**Authors:** Ruofan Yu, Hanna Sas-Nowosielska, Abhisha Sawant Dessai, Amal Oubarri, Hannah Kim, Christine L. Faunce, Ben Vanderkruk, Charly R. Good, Katherine A. Alexander, Son C. Nguyen, Ian D. Krantz, Eric F. Joyce, Melike Lakadamyali, Shelley L. Berger

## Abstract

Nuclear speckles have long been recognized as RNA-rich nuclear bodies, yet their role in genome organization and gene regulation remains incompletely understood. Using a rapid dTAG-mediated degradation system to simultaneously deplete SON and SRRM2—the core structural components of nuclear speckles—we identify a novel class of genomic regions, which we term SWING regions. Upon speckle disruption, SWING regions relocate to the nuclear periphery and acquire repressive histone marks such as H3K9me3, accompanied by gene downregulation, particularly of genes involved in developmental pathways. Consistent with this, depletion of Lamin A reduces lamina association of SWING regions and enhances their association with nuclear speckles, supporting a bidirectional balance between these two nuclear compartments. Notably, human mutations in SON and SRRM2 are associated with neurodevelopmental disorders characterized by intellectual disability and global developmental delay. Patient-derived cells bearing such mutations similarly exhibit SWING-region relocalization and gene repression, underlining a role for speckles in developmental gene regulation. We also report that drug-induced speckle rejuvenation partially rescues aberrant SWING-region localization to the nuclear lamina in patient-derived cells and iPSCs with acute depletion of SON. These findings identify SWING regions as an intermediate chromatin state positioned between nuclear speckles and the lamina, maintained by opposing functions from both structures. Our work reveals a mechanism underlying the contribution of nuclear bodies to 3D genome organization, highlights the importance of nuclear speckles and SWING regions in developmental regulation, and provides a potential therapeutic intervention in speckle dysfunction.

**Impact Statement:** Identification of nuclear speckles as determinants of specific 3D genome organization.

Demonstration of functional interactions between opposing nuclear structures (speckle versus lamina) through SWING regions.

Establishment of developmental and human disease relevance of speckle-mediated genome organization.

Providing potential avenues for therapeutic intervention in speckleopathies.

## Introduction

Nuclear bodies are membraneless, morphologically distinct sub-nuclear compartments increasingly recognized as active participants in genome function rather than passive storage depots. Among them, nuclear speckles are enriched for RNAs and RNA-processing proteins and have been implicated in both chromosome organization and gene regulation^1,2^. In parallel with broader efforts to integrate structure and function in the 4D nucleome, evidence has accumulated that speckles help coordinate transcription, splicing, 3′-end formation, and mRNA export, positioning them as dynamic hubs for gene expression^3,4^.

Genomic mapping of speckles^5–8^ showed that genomic regions closest to speckles—so-called speckle-associated domains (SPADs)—harbor dense clusters of actively transcribed genes and exhibit euchromatic features, including H3K27ac and H3K4me3 enrichment and robust RNA polymerase II occupancy; importantly, the mean distance of a gene to speckles is a strong predictor of expression across cell types^6^. Consistent imaging and genomic analyses emphasize that speckle proximity marks transcriptionally permissive neighborhoods enriched for RNA-processing machinery and active histone modifications^5–8^. In contrast, lamina-associated domains (LADs) are depleted of active histone modifications, enriched for repressive heterochromatin (for example, H3K9me3-associated states), and display low gene activity, underscoring a spatial and epigenetic opposition between speckle-proximal and lamina-proximal compartments crucial to gene expression state^9,10^. Despite this mutual exclusivity, how the compartments are stably demarcated and whether there is an active competition between them relevant to gene regulation, is not known.

Further, although there is striking correlation between SPADs and active chromatin^5–8^, mechanistic insight is lacking into the contribution of speckles to chromatin organization. Our previous work implicated chromatin architectural factors—CTCF and the cohesin subunit RAD21—in promoting speckle–chromatin association, providing evidence that speckles participate in 3D genome architecture^8^. However, relatively few perturbation studies have directly tested speckle function in active chromatin organization, and those available have generally reported only modest effects on chromatin architecture^11,12^, a confounding outcome that suggests it may be necessary to look beyond active chromatin to uncover how speckles exert broad organizational roles.

It is notable that speckles appear to have profound relevance to human health. The clinical relevance of speckle integrity is highlighted by “nuclear speckleopathies”^13^. These are inherited human variants in core speckle components^14^, including within SON^15^ and SRRM2^16^, that are associated with neurodevelopmental disorders. Severe speckleopathies are characterized by intellectual disability and global developmental delay ^13^, underscoring the importance of speckles to normal human development.

Here, we investigate a role of nuclear speckles in broad genome organization and gene expression. We acutely disrupt speckles via rapid dTAG-mediated co-degradation of SON and SRRM2, leading to identification of a speckle-wired genomic program and revealing a new class of genomic regions (“SWING” regions) defined by behavior upon speckle disruption: relocalization to the nuclear lamina, acquisition of repressive chromatin features, and gene downregulation. We further show that perturbing the nuclear lamina promotes SWING region association with speckles, revealing a competitive speckle–lamina axis that controls SWING region positioning. We find that developmental genes are particularly enriched within the SWING regions, along with broader morphogenetic and tissue-organization genes. We show that patient-derived cells bearing a SON mutation exhibit similar SWING region relocalization to the lamina and gene repression, thus correlating speckle integrity to developmental gene control in human disease. Finally, we show that pharmacologic “speckle rejuvenation” can partially rescue aberrant SWING region localization and gene expression both in such patient-derived cells and in an iPSC model with rapid SON degradation, suggesting potential for therapeutic intervention in speckle-associated pathologies.

## Results

### 1. Disruption of nuclear speckles leads to increased lamina association of certain genomic domains

The core nuclear speckle components SON and SRRM2—both necessary for speckle formation^14^ —are required for cell and organismal viability^17,18^. To enable temporal control of speckle disruption, we used a previously constructed K562 cell line in which both SON and SRRM2 were tagged with FKBP12^F36V^, allowing acute depletion with the small molecule dTAG^11^. Depletion of both proteins was evident by 24h (**Figure 1A** and **Supplementary Figure 1A**). Nuclear speckles were effectively dispersed after 24h of dTAG treatment: proteins normally enriched in speckles, including RBM25^19,20^ and SF3a66^7^, lost their speckle-like localization and redistributed throughout the nucleoplasm with reduced granularity and more uniform distribution (**Supplementary Figure 1B-C**).

**Figure 1.**
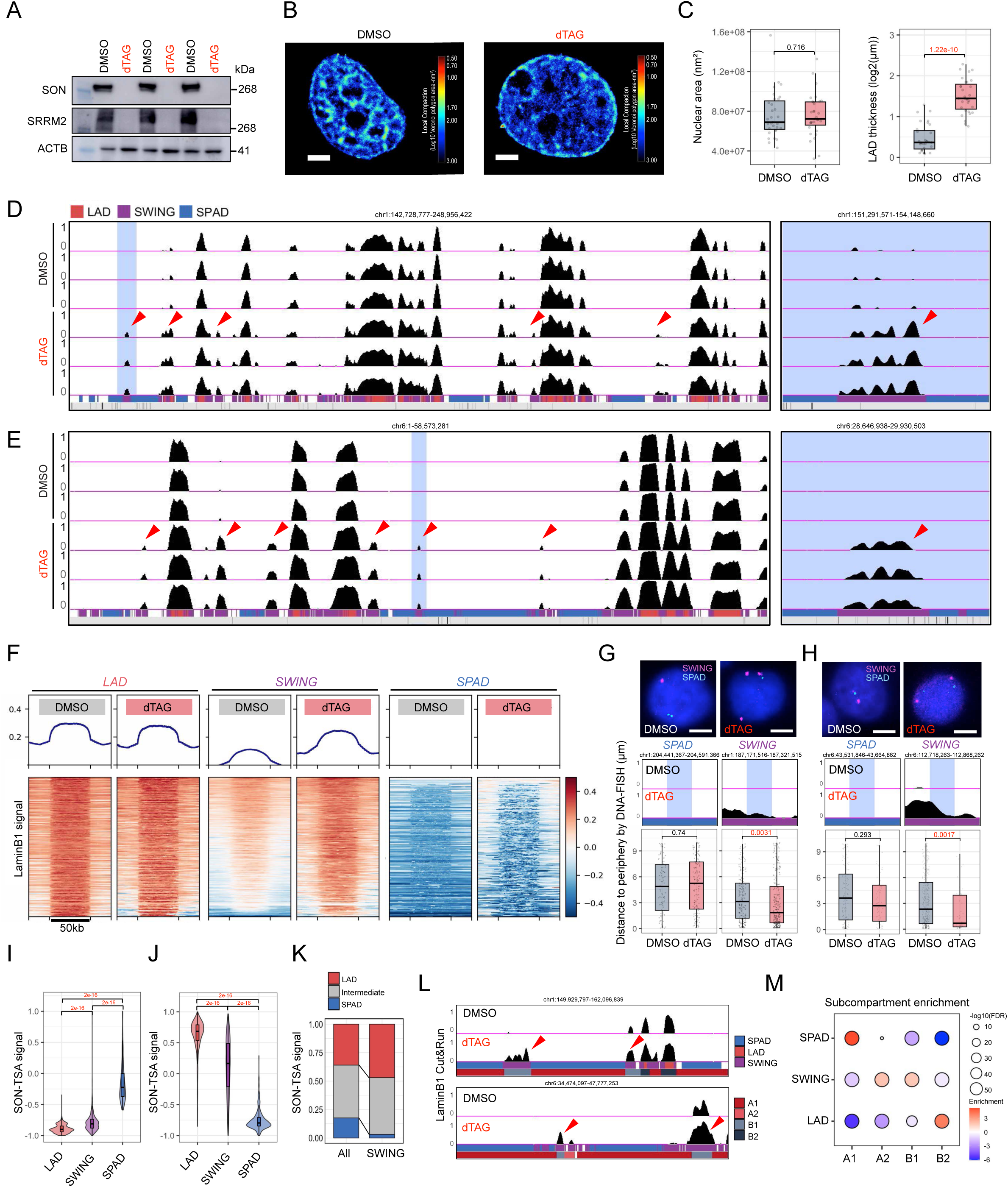
Disruption of nuclear speckles increases lamina association of specific genomic domains. **(A)** Western blot of FKBP12-tagged core speckle proteins SON and SRRM2 after 24hr of dTAG treatment; ACTB serves as a loading control. **(B)** Representative STORM image from K562 SON/SRRM2 double knock-in cells with or without dTAG treatment for 24hr; the color scale encodes log10 Voronoi polygon area. **(C)** Quantification of nuclear area (left) and LAD thickness (right) from STORM datasets. Box plots show the median and interquartile range (IQR); whiskers extend to 1.5× IQR. P-values were computed using two-sided Welch’s t-tests with Benjamini–Hochberg correction for multiple comparisons. **(D–E)** Representative genome browser tracks (three biological replicates per condition) for DMSO and dTAG-treated samples. Red arrows mark regions with increased lamina association upon dTAG treatment. Bottom tracks indicate domain calls: LAD (red), SWING (purple), and SPAD (blue). A zoomed-out view of the light-blue highlighted region is shown at right. **(F)** Metagene profiles and heatmaps of Lamin B1 CUT&RUN signal across all called LADs, SWING regions, and SPADs in DMSO versus dTAG-treated K562 cells. **(G–H)** (Top) DNA-FISH images using probes targeting a SPAD locus (cyan) and a SWING locus (magenta) in DMSO- or dTAG-treated cells. (Bottom) Corresponding Lamin B1 CUT&RUN browser tracks and quantification of 3D distances from FISH signals to the nuclear periphery. Box plots as in (C); *P* values by two-sided Mann–Whitney *U* test. Each dot represents one FISH foci. **(I–J)** Quantification of speckle association (SON TSA-seq signal; I) and nuclear lamina association (Lamin B1 DamID signal; J) across LADs, SWING regions, and SPADs in K562 cells. Violin plots show distributions with embedded box plots (median, IQR; whiskers 1.5× IQR) where applicable. *P* values are from two-sided Mann–Whitney U tests. **(K)** Distribution of lamin-associated domains (LADs), speckle-associated domains (SPADs), and intermediate regions (not classified as LAD or SPAD) in baseline K562 cells, shown genome-wide (“All”) and within SWING regions. **(L)** Representative view in K562 cells showing Lamin B1 CUT&RUN signal, corresponding SWING region calls, and Hi-C subcompartment annotations. **(M)** Enrichment of A1, A2, B1, and B2 subcompartments within LADs, SWING regions, and SPADs in K562 cells, color represents enrichment (Log2(observed/expected)) and bubble size corresponds to statistical significance (−log10 FDR).

To assess changes in chromatin organization at the single nucleosome level upon speckle disruption, we performed Stochastic Optical Reconstruction Microscopy (STORM) imaging of immunolabeled total histone H3. H3 protein levels remained unchanged following dTAG treatment to deplete SON/SRRM2 (**Supplementary Figure 1D**). However, we observed striking redistribution of H3 toward the nuclear periphery (**Figure 1B** and **Supplementary Figure 1E**). To quantify changes in chromatin organization in an unbiased fashion, we applied Objective Single Molecule Nuclear Architecture Profiler (O-SNAP)^21,22^ analysis to extract 144 features representing the spatial distribution of chromatin domains within the nucleus from the STORM H3 images. This analysis revealed significant changes to chromatin domain organization at the nuclear periphery (see **Supplementary Figure 1E** for all imaged nuclei and **Supplementary Figure 1F** for summary of O-SNAP significant features). One prominent change was significant increase of the thickness of Lamin Associated Domains (LADs) modeled from the STORM images of chromatin at the nuclear periphery (**Figure 1C, right**). This analysis revealed increased association of chromatin with nuclear lamina, with no apparent change in nuclear size (**Figure 1C, left**).

In light of this observation—that speckle disruption relocates chromatin to the lamina—we investigated whether the effect is global or confined to specific genomic regions, via assaying DNA association with the lamina utilizing Lamin B1 CUT&RUN^23^. Our Lamin B1 CUT&RUN profiles showed strong technical reproducibility (**Supplementary Figure 2A**) and correlated well with previous Lamin B1 DamID^24^, a widely used method for mapping lamina-associated regions (Supplementary Figure 2B). We also note that Lamin B1 subcellular localization and protein levels were not visibly altered in 24h dTAG-treated cells (**Supplementary Figure 2C-D**). Upon 24h of dTAG treatment, we observed striking gains of Lamin B1 association with specific DNA regions, rather than a global increase (**Figure 1D-E**). These regions comprise the subset of genomic loci that specifically acquire lamina association upon speckle disruption, distinguishing them from constitutive LADs and regions that remain lamina-detached. Because they undergo this inducible shift in lamina association, we refer to them as “SWING” domains to highlight their dynamic behavior (see purple regions in the BED tracks in **Figure 1D–E**). Consistently, the metagene plot and heat map show a clear increase in lamina association, represented by an increase of Lamin B1 CUT&RUN signal within SWING regions, but not within canonical LADs nor within speckle-associated domains (SPADs) (**Figure 1F**). To further substantiate this conclusion, we performed DNA-fluorescence in situ hybridization (FISH) to measure the distance between probe foci and the nuclear periphery. In agreement with the Lamin B1 CUT&RUN results, SWING probes relocated significantly closer to the lamina, whereas SPAD probes showed no significant movement (**Figure 1G-H**). This lack of SPAD movement may reflect the presence of additional stabilizing features of active DNA regions—such as CTCF/cohesin loops^8^ and active transcriptional machinery^5^ —that may help to maintain their nuclear position distant from the lamina even after 24 h of speckle disruption. In summary, we observed that a subfraction of the genome (30.45%, see **Supplementary Figure 2E** for distribution)—here newly named “SWING” domains—gain nuclear lamina association upon acute disruption of nuclear speckles.

### 2. SWING regions exhibit intermediate properties

We next investigated certain properties of SWING regions. Using previously published genomic maps of speckles (SON TSA-seq) and the nuclear lamina (Lamin B1 DamID) in K562 cells^6,24^, we found that SWING regions exhibit intermediate association levels with both structures (**Figure 1I–J, Supplementary Figure 3A**), indicating SWING is located between speckles and nuclear lamina. To assess whether this conclusion holds cytologically, we analyzed genome-wide multiplex FISH (MERFISH) data generated in IMR90 cells^25^. Although this is a different cell line, prior work reported some conservation of both SPADs and LADs across cell lines^6,10,26^. Consistent with the genomic profiles, SWING regions in the MERFISH dataset occupied intermediate distances to both speckles and the lamina (**Supplementary Figure 3B–D**), and slightly trending more towards lamina, suggesting that SWING regions may occupy a more repressed-leaning position than SPAD. Supporting this notion, we examined the baseline localization of SWING regions in K562 cells prior to dTAG-mediated speckle disruption and found that (more than 96%) localize either to LADs or to intermediate regions not classified as LAD or SPAD, rather than to SPADs (**Figure 1K**; bars represent the fraction of genomic regions in each category across chromosomes; “All” denotes the genome-wide distribution of LAD, SPAD, and intermediate regions; single-chromosome view in **Supplementary Figure 3E**).

At a broader genomic scale, lamina association shows partial conservation across cell lines, with a subset of lamina-associated domains (LADs) shared across cell types while other LADs are more dynamic^26^. Given the SWING region sensitivity to speckle perturbation and its intermediate nature, we investigated whether the SWING region is variable across cell lines. We compared published Lamin B1 association profiles^24^ for K562 (chronic myeloid leukemia), HFFc6 (human foreskin fibroblast), HCT116 (colorectal cancer), and H1 (human embryonic stem cell) cell lines. We calculated a variability score by quantifying the standard deviation of the Lamin B1 signal across these four cell lines and categorized them as low, medium, or high variability. Unlike canonical LADs which have relatively consistent Lamin B1 distribution across cell lines, in SWING regions Lamin B1 association exhibited significant variability between cell lines, where 42% of SWING regions were associated with medium- to high-variability Lamin B1 regions (**Supplementary Figure 3F–H**; see red-shaded canonical LADs and purple-shaded SWING regions for comparison.). Together, these observations suggest that a substantial subset of regions traditionally annotated as LADs may behave functionally as SWING regions—displaying intermediate characteristics and heightened sensitivity to perturbation of nuclear speckle.

We next asked whether SWING regions also display intermediate chromatin architecture. Because SPADs are strongly associated with the active A1 subcompartment^5,27^ and LADs with the repressive B2/B3 subcompartment^27^, we examined subcompartment composition within SWING regions. Using published Hi-C data from K562 cells^28^, we assigned genomic subcompartments^29^ and observed that SWING regions frequently co-occur with the intermediate A2 and B1 subcompartments, and thus not strongly associated with activation or repression (**Figure 1L**; see upper BED track for LAD, SWING and SPAD localization and bottom track for subcompartment annotation). Genome-wide analyses further revealed that SWING regions are enriched for A2 and B1, whereas both SPADs and LADs are depleted for these intermediate subcompartments (**Figure 1M, Supplementary Figure 3I**). Thus, even when considering chromatin partitioning alone, independent of lamina or speckle association, SWING regions exhibit intermediate properties that are distinct from LADs and SPADs, suggesting a unique biological role.

### 3. SWING region becomes more heterochromatin-like upon speckle disruption

Given that SWING regions show intermediate association with the nuclear lamina and nuclear speckles, we next investigated whether they also display hybrid histone modification profiles, between activation-associated acetylation and repression-associated methylation. Using publicly available ENCODE ChIP–seq datasets^30^ from K562 cells, we quantified the distribution of six histone modifications. For all six modifications, SWING regions showed signal levels between those observed in LADs and SPADs (**Figure 2A**). Notably, SWING regions exhibited intermediate enrichment for the facultative heterochromatin mark H3K27me3 (high in SPADs) and, in particular, for the constitutive heterochromatin mark H3K9me3 (high in LADs) (**Figure 2A**). These two modifications appear in a heterogeneous pattern across SWING regions—some regions enriched for H3K27me3 and others for H3K9me3—in contrast to the more uniform H3K27me3 in SPADs and H3K9me3 in LADs (sample region in **Figure 2B**). Together, these patterns indicate that SWING regions represent an intermediate, relatively repressed chromatin state that bridges constitutive and facultative heterochromatin and shares features of both.

**Figure 2.**
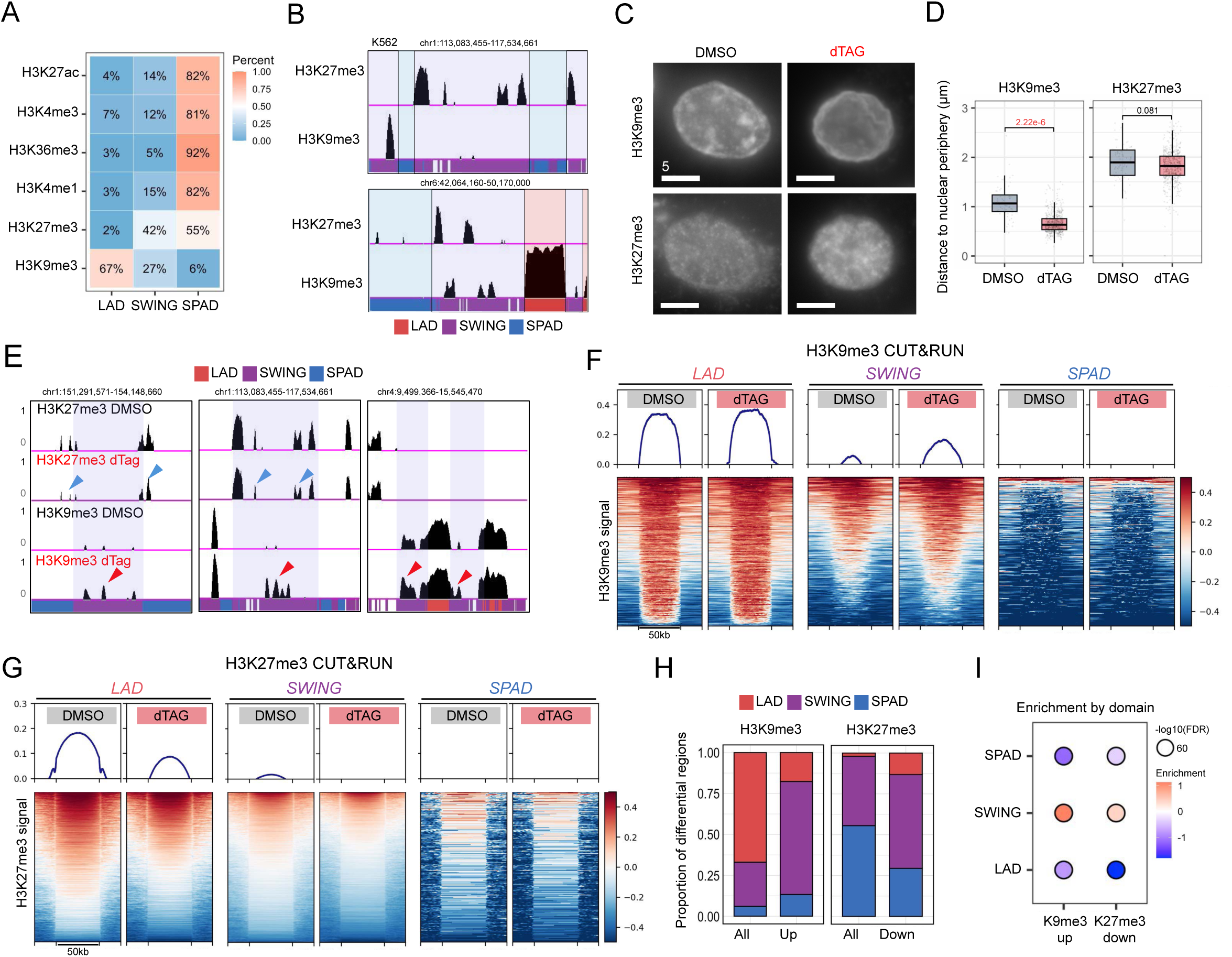
SWING regions become more heterochromatin-like upon speckle disruption. **(A)** Percentile distributions of designated histone-modification ChIP–seq signals across LADs, SWING regions, and SPADs. **(B)** Example browser view of H3K27me3 and H3K9me3 CUT&RUN signal in control K562 cells. BED tracks indicate domain calls: LAD (red), SWING (purple), SPAD (blue). **(C)** Representative IF images for H3K9me3 and H3K27me3 in K562 cells with or without 24 h dTAG treatment. **(D)** Quantification of distances from the nuclear periphery for regions classified as H3K9me3 or H3K27me3 in K562 cells with or without 24 h dTAG treatment. Box plots show the median and interquartile range (IQR); whiskers extend to 1.5× IQR. P values by two-sided Mann–Whitney U tests. **(E)** Example browser view of H3K27me3 and H3K9me3 CUT&RUN signal in K562 cells with or without 24 h dTAG treatment; blue arrows indicate regions with reduced signal and red arrows indicate regions with increased signal. **(F–G)** Metagene profiles and heatmaps of H3K9me3 (F) and H3K27me3 (G) CUT&RUN signal across all called LADs, SWING, and SPADs in DMSO versus dTAG-treated K562 cells. Metagene lines show mean normalized CUT&RUN signal with shaded 95% CI; heatmaps display z-scored signal per region, centered at region midpoints and ordered by mean signal. **(H)** Percentages of (left) all differential regions and up-regulated H3K9me3 regions and (right) all differential regions and down-regulated H3K27me3 regions within LADs, SWING regions, and SPADs. Bars show category proportions. **(I)** Enrichment of H3K9me3-upregulated regions and H3K27me3-downregulated regions across LADs, SWING, and SPADs, reported as log2 (observed/expected).

We next tested whether acute speckle disruption alters H3K9me3 and H3K27me3 distribution in the nucleus, and in particular within SWING regions. Immunofluorescence in cells with or without dTAG-mediated speckle disruption showed that H3K9me3 underwent striking redistribution toward the nuclear periphery, whereas H3K27me3 remained broadly evenly dispersed across the nucleus (**Figure 2C**; quantification in **Figure 2D**). Because SWING regions relocalize to the nuclear lamina—a nuclear context associated with H3K9me3 deposition—we examined whether these changes occur specifically within SWING. We utilized CUT&RUN profiling for H3K9me3 and H3K27me3 and focused on the SWING regions previously defined (see **Figure 1**). We observed a clearly pronounced increase in H3K9me3 that was specific to SWING (**Figure 2E** and **Figure 2F**). Consistent with this, most regions with significantly gained H3K9me3 upon speckle disruption mapped to SWING (**Figure 2H**, left), showing a clear enrichment (**Figure 2I**). In contrast, H3K27me3 showed a decrease at both SWING and LAD (example tracks in **Figure 2E**; genome-wide quantification in **Figure 2G**), with depletion across SWING that was weaker than the gain observed for H3K9me3 (**Figure 2H**, right; **Figure 2I**). Together, the increased H3K9me3 and reduced H3K27me3 indicate that, upon acute speckle disruption, SWING chromatin shifts towards a more constitutive heterochromatin-like state.

### 4. Nuclear Speckle Disruption leads to down-regulation of SWING genes

We next investigated whether the increased lamina association and gain of repressive H3K9me3 in SWING regions upon acute speckle depletion led to changes in gene expression. We performed RNA-seq in K562 cells with 24h of dTAG treatment to disrupt nuclear speckles, or with DMSO control. As quality controls, we observed a general shift toward gene downregulation in dTAG-treated samples (**Supplementary Figure 4A**). We validated that LADs, SWING, and SPADs show progressively increased numbers of expressed genes and increasing basal expression (**Supplementary Figure 4B**), consistent with previous observations that gene expression increases with speckle association and decreases with lamina association^5^. Consistent with prior reports that speckles promote expression of speckle-associated genes^7,8,31,32^, genes in SPADs were predominantly downregulated upon speckle disruption (**Figure 3A-B**). Strikingly, genes in SWING regions were also significantly downregulated, and, notably, to a greater extent than were SPAD genes in magnitude of fold-change (**Figure 3A-B**). This difference was not explained by baseline expression: in an expression-matched analysis, SWING genes still showed a stronger tendency toward downregulation than SPAD genes (**Figure 3C**). Unsupervised hierarchical clustering revealed that SWING-region genes segregate into a prominent cluster within the differentially expressed gene set that clearly separates from SPAD genes (**Figure 3D**). Although more SPAD genes change overall (824 SPAD vs. 359 SWING), reflecting the higher gene density of SPAD regions, the magnitude and direction of change differ sharply: SWING genes show a pronounced bias toward downregulation, as evident from both the volcano plot and the density distribution (**Figure 3E**). Genome browser views of top LaminB1 differential genes further illustrate these trends: genes within SWING regions showed increasing Lamin B1 CUT&RUN signal and tendency to be downregulated upon speckle disruption (**Figure 3F**). Of note, because very few constitutive LAD (defined as FDR > 0.1 in Lamin B1 C&R in DMSO vs. dTAG) genes are expressed in these cells, they were not included in this comparison.

**Figure 3.**
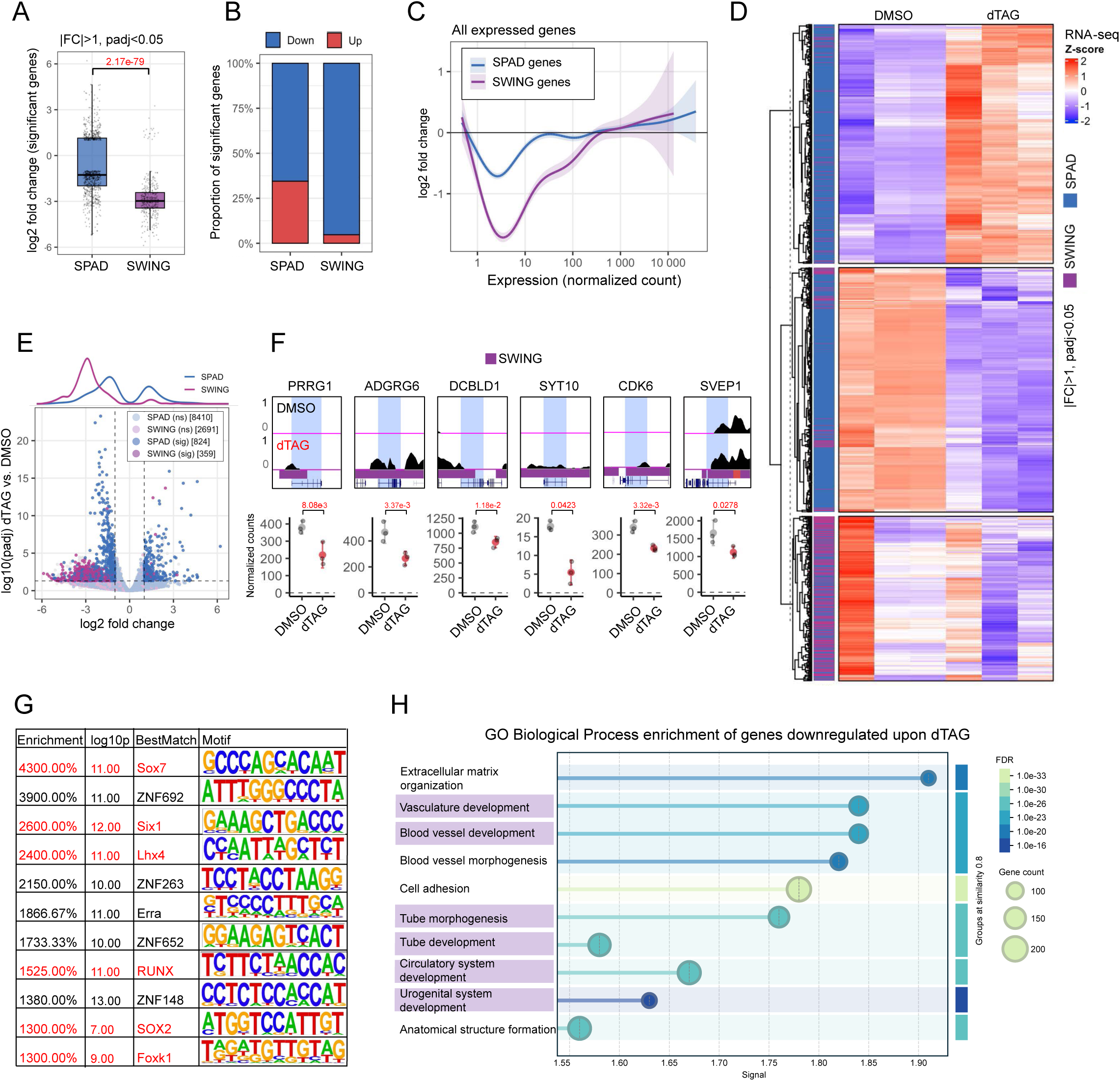
Nuclear speckle disruption leads to down-regulation of SWING genes. **(A)** Comparison of significantly differentially expressed genes (FDR < 0.05 and |log2 fold change| > 1) between dTAG 24 h and DMSO in K562 cells, grouped by localization (SPAD vs SWING). Box plots show median and IQR; whiskers, 1.5× IQR. P values by two-sided Mann–Whitney U tests. **(B)** Percentile distributions showing the proportions of significantly down-regulated (blue) and up-regulated (red) genes within SPAD or SWING. **(C)** Distributions of log2 fold change for expression-matched SPAD versus SWING genes (no FDR or fold-change cutoff). Lines show mean with shaded 95% CI. **(D)** RNA-seq heatmap of all significantly differentially expressed genes (FDR < 0.05 and |log2 fold change| > 1) in SWING and SPAD with hierarchical clustering. The y-axis lists individual genes; SPAD genes are labeled blue and SWING genes purple. Values are z-scores of normalized RNA-seq counts. **(E)** Volcano plot for K562 cells treated with dTAG (24 h) versus DMSO. Points are colored by domain localization (SPAD, blue; SWING, purple). “Sig” indicates significantly differentially expressed genes (FDR < 0.05 and |log2 fold change| > 1); “NS” indicates not significant. The panel above shows density distributions for the designated gene groups. **(F)** Example browser view showing Lamin B1 CUT&RUN tracks and normalized RNA-seq counts for designated genes within SWING regions. **(G)** HOMER motif-enrichment results for SWING regions; entries highlighted in red denote development-related transcription factors. **(H)** Gene Ontology Biological Process enrichment for significantly downregulated genes (FDR < 0.05 and log2 fold change < −1) in dTAG 24 h versus DMSO K562 cells; development-related terms are highlighted.

Beyond the well-documented role of speckles in promoting expression of SPAD genes, our data reveal that speckles also regulate genes in SWING regions, which lie at intermediate distances from speckles. Because SPAD genes are enriched in cellular processes requiring high basal and/or rapid expression^6,33^ (e.g., housekeeping and stress-response pathways), we investigated whether genes within SWING regions also encompass preferential pathway associations. Motif analysis across all identified SWING regions revealed that top-ranked transcription factors enriched include developmental regulators, such as the embryonic and cell-fate determinant SOX7^34^ and the homeobox factor SIX1^35^ (**Figure 3G**, developmental TFs highlighted. Consistently, gene ontology analysis of SWING-localized genes showed significant enrichment for development-related pathways (**Supplementary Figure 4C**). In addition, genes downregulated upon acute speckle disruption were significantly enriched for development-associated morphogenetic and tissue-organization programs (**Figure 3H**). Together, these results suggest that, in contrast to SPAD genes that require high basal and/or rapid expression, SWING genes are subject to developmentally induced activation or repression and are particularly sensitive to speckle perturbation.

### 5. SWING regions lose lamina association upon LMNA disruption

Building on our finding that SWING regions relocalize toward the nuclear lamina after speckle disruption—accompanied by increased H3K9me3 and gene downregulation—we investigated whether speckles and the lamina might actually “compete” for association at SWING regions. Prior studies suggest that speckle- and lamina-associated chromatin are largely mutually exclusive^24,27^, raising a question of whether “swinging” occurs bidirectionally, in particular, whether weakening the nuclear lamina tethering relocates SWING regions away from the lamina and potentially toward speckles.

Chromatin–lamina association is mediated by several tethering mechanisms, one of which is through Lamin A/C (LMNA)^36^. LMNA has been strongly implicated in development and differentiation^37,38^ ; given this development relevance, we asked how LMNA loss affects SWING region localization by reanalyzing published Lamin B1 DamID datasets from K562 cells bearing CRISPR-mediated LMNA^24^ knockout (see **Supplementary Figure 5A** for Western blot validation performed in this study), using the LAD/SWING/SPAD annotations defined in our double dTAG system. LMNA knockout produced a selective reduction of lamina association in SWING regions, with minimal change across canonical LADs or SPADs (**Figure 4A**). Using the same cell lines, we performed Cut&Run against SON, a method previously validated to capture chromatin–speckle associations^8,23^. Comparison of our SON Cut&Run data with published SON TSA-seq in wild-type K562^6^ cells showed strong correlation (**Supplementary Figure 5B–C**). Upon LMNA loss, we observed a global increase in SON association (**Figure 4B**) with the most pronounced changes—both Lamin B1 loss and SON gain—occurring in SWING regions (**Figure 4C–D**). Reciprocally, differential analysis further confirmed that regions with altered Lamin B1 association in LMNA KO cells predominantly map to SWING regions (**Figure 4E**). When jointly analyzing speckle and lamina association across all identified SPAD, SWING, and LAD regions, SWING regions formed a distinct cluster characterized by Lamin B1 loss and SON gain (**Figure 4F**). These findings support our model that SWING regions are regulated distinctly from SPADs and LADs and are uniquely sensitive to perturbations in nuclear architecture.

**Figure 4.**
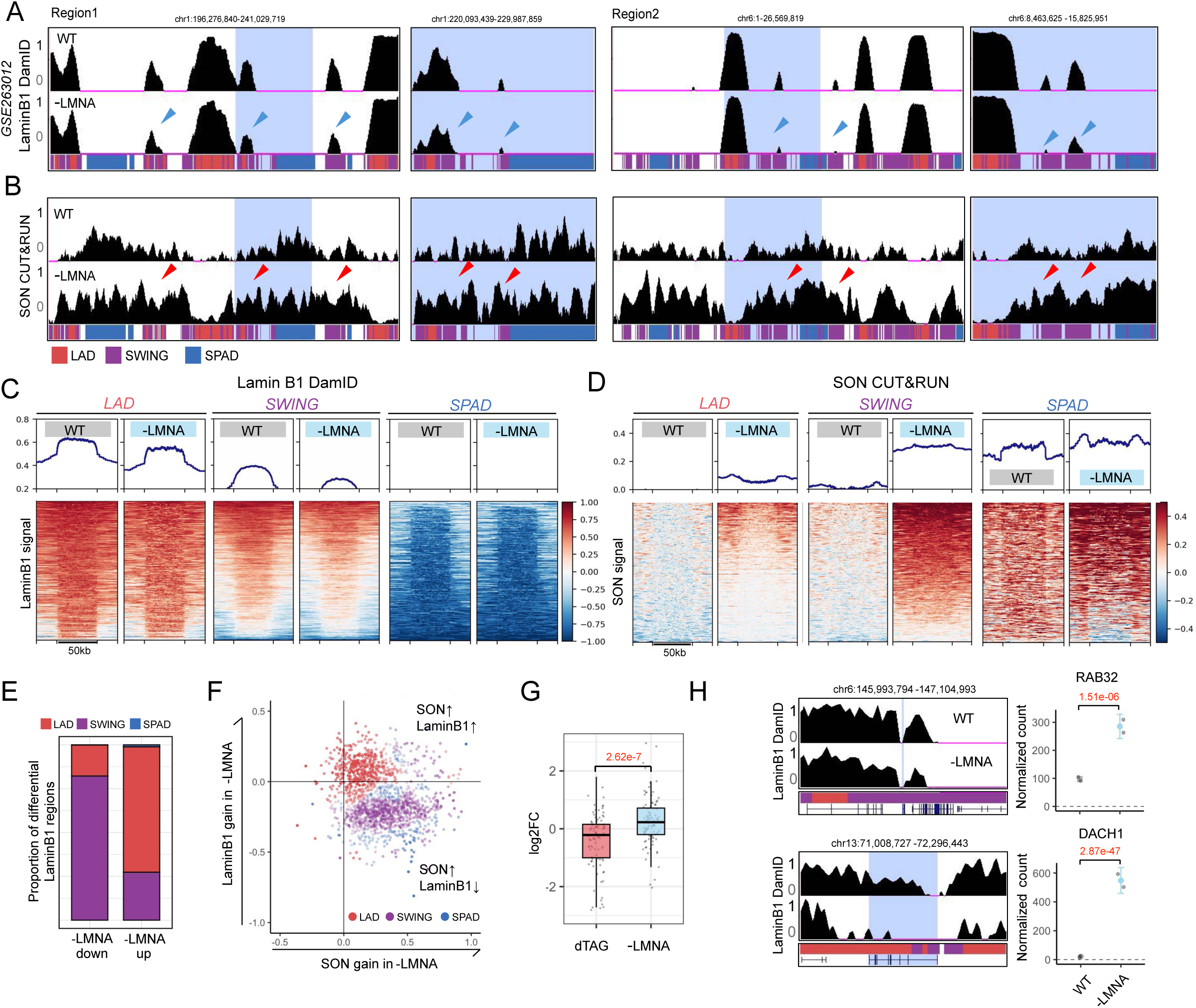
SWING regions relocate away from the lamina upon LMNA disruption. **(A-B)** Representative genome browser view of (A) Lamin B1 DamID and (B) SON Cut&Run signal in WT and LMNA KO K562 cells. Blue arrows mark regions with decreased lamina association, and red arrows mark regions with increased speckle association in LMNA KO. Bottom tracks indicate domain annotations: LAD (red), SWING (purple), and SPAD (blue). A zoomed-out view of the light-blue highlighted region is shown at right. **(C-D)** Metagene profiles and corresponding heatmaps of Lamin B1 DamID (C) and SON (D) signal across all LADs, SWING regions, and SPADs identified in WT and LMNA KO K562 cells. (E) Proportions (%) of genomic regions with significantly increased or decreased Lamin B1 association in LMNA KO relative to WT (right), colored by domain category (LAD, red; SWING, purple; SPAD, blue). Bars indicate the proportion of regions in each category. (F) Two-dimensional scatter plot showing changes in SON and Lamin B1 association all LAD, SWING, and SPAD domains identified in K562 cells. Each point represents a genomic region plotted by ΔSON (x-axis) and ΔLMNB1 (y-axis). Points are colored by density-based domain assignment (SPAD, SWING, or LAD), with color intensity reflecting assignment confidence. (G) Comparison of gene expression fold change (LMNA KO vs. WT) for all genes and genes located within regions showing significant loss of Lamin B1 association in LMNA KO relative to WT (FDR < 0.01). Box plots show median and interquartile range (IQR); whiskers extend to 1.5 × IQR. P values were calculated using two-sided Mann–Whitney U tests. (H) Example browser view showing Lamin B1 DamID tracks and normalized RNA-seq counts for indicated genes; light-blue shading highlights gene bodies.

We next investigated whether these localization changes are associated with altered gene expression profiles through our analysis of previously published datasets^24^. Consistent with the genomic localization shift, genes located in regions that show a significant decrease in Lamin B1 association in LMNA reduction were downregulated upon speckle disruption and significantly upregulated only in the LMNA KO (**Figure 4G**; see **Figure 4H** for representative Lamin B1 DamID and RNA-seq coverage of example genes). These inverse effects support a competitive model in which speckles and LMNA establish opposite regulatory influences on SWING-region genes.

In summary, our data support a bidirectional “shifting” of SWING regions between speckles and the lamina: speckle disruption relocates SWING regions toward the lamina, leading to gene repression, whereas LMNA loss relocates SWING regions away from the lamina, leading to gene de-repression.

### 6. Speckle activation by drug treatment reduces lamina tethering

Hence, loss of lamina tethering, specifically via LMNA disruption, relocalizes SWING regions away from the nuclear lamina. We investigated whether a speckle gain-of-function would produce a similar outcome. A recent study identified the FDA-approved small molecule pyrvinium pamoate (PP) as a potential speckle gain-of-function treatment. PP increases speckle volume and mimics the transcriptional response of SON overexpression by lowering the surface tension of an intrinsically disordered region in SON^39^. PP leads to robust speckle enlargement both in mouse embryonic fibroblasts and in human iPSC-derived neurons^39^. We tested the effect of PP in K562 cells and found that PP treatment markedly increased individual speckle size and speckle number using either immunofluorescence of SON or of the speckle-localized factor RBM25 IF (**Figure 5A**). Further, using STORM imaging, we observed that cells treated with 100 nM PP exhibited a slight, non-significant increase in nuclear size, while showing a significant reduction in LAD thickness (**Figure 5B-5C**, see **Supplementary Figure 6A** for all imaged nuclei and **Supplementary 6B** for summary of O-SNAP significant features). Notably, the latter LAD reduction was opposite of the LAD thickening we observed by STORM upon dTAG-mediated speckle disruption (see **Figure 1B-1C**).

**Figure 5.**
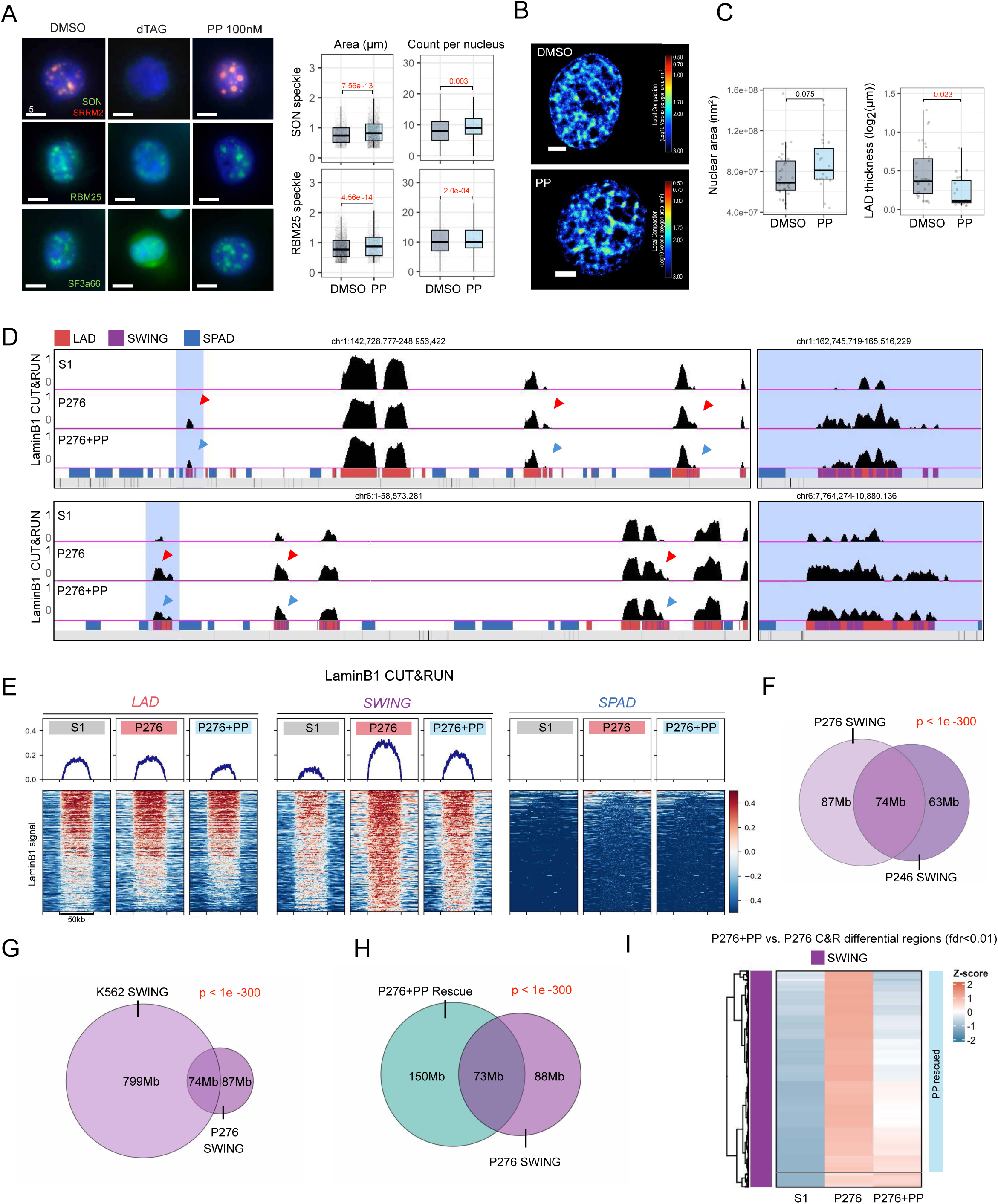
Speckle activation by drug treatment reduces aberrant lamina tethering in patient derived cells. **(A)** Left: representative IF images in K562 cells under the indicated treatments and antibodies. Right: quantification of speckle area and speckle count per nucleus. Box plots show median and IQR; whiskers, 1.5× IQR. P values by two-sided Mann–Whitney U tests. **(B)** Representative STORM image from K562 cells with or without PP (pyrvinium pamoate) treatment; the color scale encodes log10 Voronoi polygon area. **(C)** Quantification of nuclear area (left) and LAD thickness (right) from STORM datasets. Box plots as in (A). P values by two-sided Welch’s t-tests with Benjamini–Hochberg correction for multiple comparisons. **(D)** Representative genome browser view of Lamin B1 CUT&RUN signal in healthy donor (S1), patient 276 (P276), and P276 treated with 100 nM PP. Blue arrows mark regions with decreased lamina association; red arrows mark regions with increased lamina association in the indicated sample. Bottom tracks indicate domain calls: LAD (red), SWING (purple), SPAD (blue). A zoomed-out view of the light-blue highlighted region is shown at right. **(E)** Metagene profiles and heatmaps of Lamin B1 CUT&RUN signal across all called LADs, SWING regions, and SPADs, in S1, P276, and P276 with 100 nM PP. Metagene lines show mean normalized signal; heatmaps display z-scored signal per region, centered at region midpoints and ordered by mean signal. **(F)** Overlap between SWING regions identified in patient 246 and in patient 276. P value by hypergeometric test using the hg38 mappable genome as background. **(G)** Overlap between SWING regions identified in K562 and in patient 276. P value by hypergeometric test using the hg38 mappable genome as background. **(H)** Overlap between SWING regions and regions showing significant rescue of aberrant Lamin B1 under PP treatment in patient 276. P value by hypergeometric test using the hg38 mappable genome as background. **(I)** Heatmap of z-score–normalized Lamin B1 signal across all P276+PP versus P276 differential regions (FDR < 0.01) in the indicated samples, with hierarchical clustering.

Mutations in the speckle core proteins SON and SRRM2 are associated with neurodevelopmental disorders. SON mutations cause Zhu–Tokita–Takenouchi–Kim (ZTTK) syndrome^15^, and SRRM2 mutations are linked to an unnamed developmental disorder^16^. Affected individuals show overlapping phenotypes, including developmental delay, distinctive facial features, and intellectual disability, supporting a hypothesis that speckle dysfunction can lead to a group of neurodevelopmental pathologies, or “nuclear speckleopathies”^13^. For SON/ZTTK syndrome, most documented patient variants are frameshift or nonsense mutations that introduce premature stop codons; these transcripts may undergo nonsense-mediated decay, resulting in SON haploinsufficiency rather than stable truncated protein^40,41^. In contrast, much less is known about SRRM2 pathogenic mechanisms, although reported patient alleles are also predominantly premature-stop variants^16^, suggesting a similar loss-of-function model.

Our findings above reveal that speckles regulate SWING regions which are enriched for developmental pathways. Thus, we next investigated: (1) whether cells derived from patients with mutations in speckle genes exhibit aberrant gains in lamina association, and (2) whether pharmacological speckle activation using PP can restore speckle morphology and function in these patient-derived cells. We analyzed blood-derived lymphoblastoid cell lines (LCLs) from two patients: one carrying a premature stop-gain mutation in SON (P276; c.3334C>T; Sanger validation in **Supplementary Figure 7A**, patient clinical phenotype (HPO terms) in **Supplementary Figure 7B**), and the other carrying a premature stop-gain mutation in SRRM2 (P246; c.1117C>T; Sanger validation in **Supplementary Figure 7C**; HPO terms in **Supplementary Figure 7D**). Consistent with the genotype, SON expression in P276 was ∼50% of control by RNA-seq (**Supplementary Figure 7E**), and SRRM2 expression in P246 was similarly ∼50% of control (**Supplementary Figure 7F**). Immunofluorescence revealed smaller nuclear speckle area in both P246 and P276 compared with a healthy donor LCL (S1) (**Supplementary Figure 7G**; quantification in **Supplementary Figure 7H**). We then treated cells with PP, predicting that its reported functional specificity toward SON would preferentially restore speckle organization in SON-mutant cells but not in SRRM2-mutant cells. Consistent with this prediction, speckle area was restored in P276 (**Supplementary Figure 7H**; S1 vs. P276+PP), whereas no significant restoration was observed in P246 (**Supplementary Figure 7H**; S1 vs. P246+PP), supporting PP’s functional specificity toward SON^39^.

We performed Lamin B1 CUT&RUN in the patient cell lines. Consistent with our K562 dTAG results of depletion of speckles (see **Figure 1**), both P276 (SON mutation; browser view in **Figure 5D**; genome-wide summary in **Figure 5E**) and P246 (SRRM2 mutation; browser view in **Supplementary Figure 8A**; genome-wide summary in **Supplementary Figure 8B**) showed aberrant increases in lamina association specifically within SWING regions, with little or no change across canonical LADs or SPADs. Approximately 50% of SWING regions identified in the two patient lines overlap (**Figure 5F**), suggesting that SON and SRRM2 regulate chromatin positioning through an overlapping mechanism. SWING regions identified in P276 significantly overlapped with those defined in K562 (**Figure 5G**) and were similarly highly enriched for nervous system development pathway genes (**Supplementary Figure 8C**).

Notably, 100 nM PP treatment partially reduced the aberrant Lamin B1 association in P276 (example locus in **Figure 5D**; genome-wide summary in **Figure 5E**). Across all SWING regions in P276, nearly half were rescued by PP (**Figure 5H**), and reciprocally, the vast majority of differential regions within SWING were restored upon PP treatment (**Figure 5I**). We also note that, consistent with the inability of PP to rescue the reduced speckle area in P246 as assessed by immunofluorescence (**Supplementary Figure 7G-H**), although aberrant lamina association was apparent in P246 carrying an SRRM2 mutation, a PP-dependent reduction in Lamin B1 signal was not readily detectable (**Supplementary Figure 8A–B**). These observations suggest that the efficacy of PP operates specifically through SON, as previously reported^39^.

In summary, patient cells harboring disease-causing mutations in SON or SRRM2 exhibit abnormal increases in chromatin–lamina associations; in one patient bearing mutation in SON (P276), this phenotype was ameliorated by PP treatment, consistent with speckle activation restoring both speckle morphology and lamina association toward a healthy state.

### 7. Developmental pathway dysregulation occurs in speckleopathy patient cells

We next investigated whether the aberrant lamina association observed in the patient lines—and partial reversal by PP—extends to gene expression changes. We performed RNA-seq in LCLs from the healthy donor (S1) and from P276 and P246. For P276, genes located within SWING regions (the regions showing aberrant lamina gain) were overwhelmingly downregulated relative to S1 (distribution in **Figure 6A**; domain-focused comparison in **Figure 6B**). Compared with SPAD or LAD genes (of which about half were significantly downregulated versus upregulated) more than 90% of differentially expressed SWING genes were downregulated (**Figure 6C**; heatmap in **Figure 6D**). These data mirror our acute speckle disruption results in K562 dTAG cells (see **Figure 3**).

**Figure 6.**
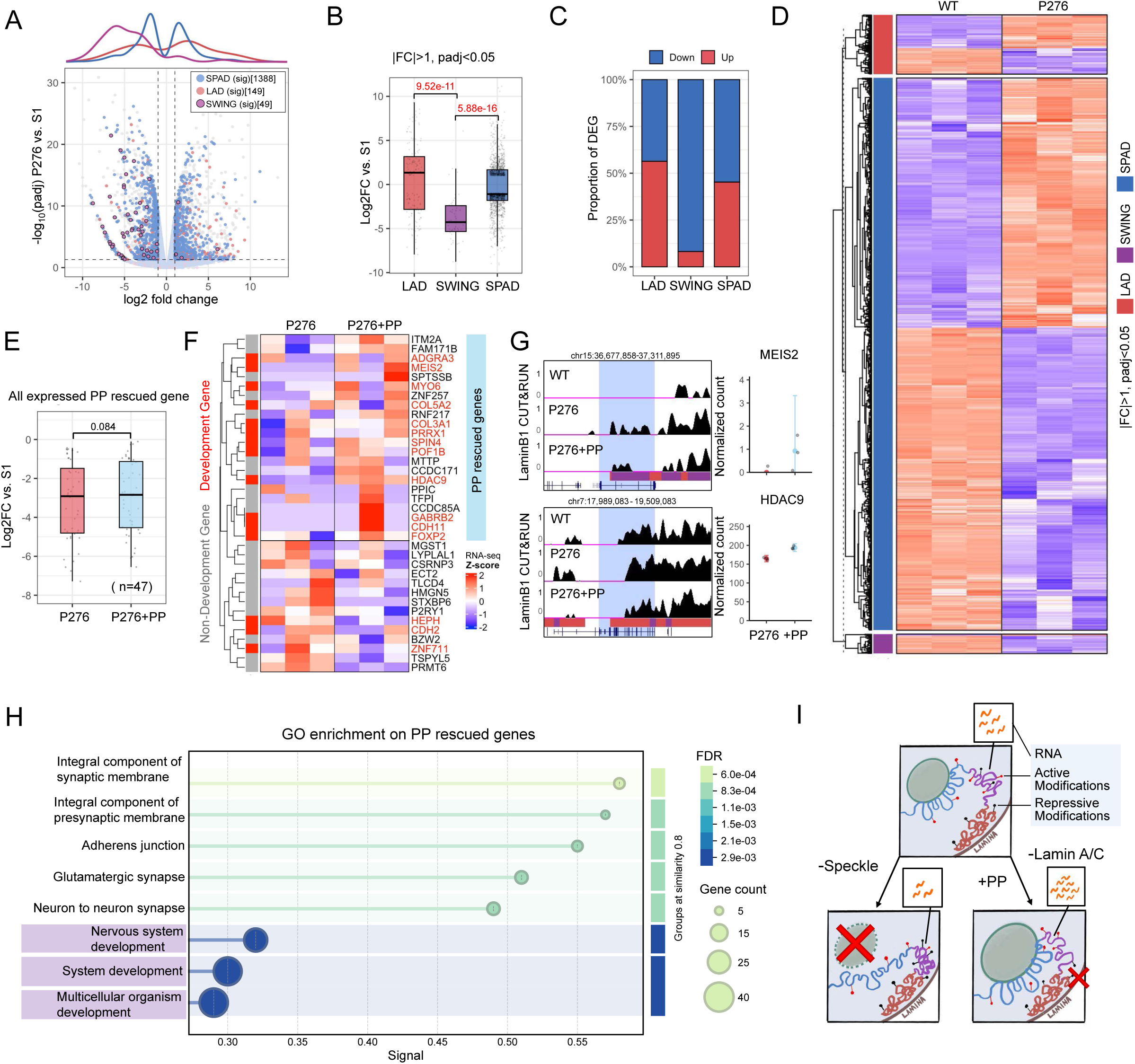
Developmental pathway dysregulation in speckleopathy patient cells is mitigated by pyrvinium pamoate. **(A)** Volcano plot of RNA-seq differential analysis between patient 276 (P276) and healthy donor (S1). Points are colored by domain localization (SPAD, blue; SWING, purple; LAD, red). “Sig” denotes significantly differentially expressed genes (FDR < 0.05 and |log2 fold change| > 1); “NS” denotes not significant. Axes: x, log2 fold change; y, −log10 FDR. The panel above shows density distributions for the designated gene groups. **(B)** Comparison of significantly differentially expressed genes (FDR < 0.05 and |log2 fold change| > 1) between P276 and S1, grouped by localization (SPAD, SWING and LAD). Box plots show median and IQR; whiskers, 1.5× IQR. P values by two-sided Mann–Whitney U tests. **(C)** Percentile distributions showing the proportions of significantly down-regulated (blue) and up-regulated (red) genes within SPAD, SWING or LAD. **(D)** RNA-seq heatmap of all significantly differentially expressed genes in SWING, SPAD and LAD with hierarchical clustering between P276 and S1. The y-axis lists individual genes; Values are z-scores of normalized RNA-seq counts (per-gene z-scaling; distance = 1 − Pearson correlation; average linkage). **(E)** Comparison of log2 fold change relative to S1 for genes that show Lamin B1 rescue under PP (100 nM) in P276 (untreated) versus P276+PP. P values by two-sided Wilcoxon signed-rank test. **(F)** RNA-seq heatmap of genes with Lamin B1 rescue under PP that are significantly affected by PP (FDR < 0.10); development-related genes are highlighted in red. **(G)** Example browser view showing Lamin B1 CUT&RUN tracks and normalized RNA-seq counts for indicated genes; light-blue shading highlights gene bodies. **(H)** Gene Ontology Biological Process enrichment for genes rescued by PP (RNA-seq: P276+PP vs P276, FDR < 0.10 and fold change > 0; Lamin B1 CUT&RUN: decreased Lamin B1 in P276+PP vs P276, FDR < 0.01). Development-related terms are highlighted. **(I)** Schematic illustrating the model of joint regulation of SWING regions by nuclear speckles and nuclear lamina.

For P246, we observed a similar pattern: differentially expressed SWING genes were strongly downregulated relative to S1 (**Supplementary Figure 9A–B**), whereas SPAD and LAD genes showed a roughly balanced mix of down- and upregulation (**Supplementary Figure 9C**). Together, these findings indicate that SWING genes are highly sensitive to speckle-mediated control in a patient context with long-term speckle dysfunction.

We next assessed whether PP treatment modulates gene expression at loci with aberrant lamina gain that were rescued at the chromatin level. Given that LCLs are non-differentiating cell lines, baseline expression of developmental genes is low, and we therefore anticipated that transcriptional rescue, particularly for neurodevelopmental genes, may be modest and more difficult to detect. Because robust PP-mediated rescue was observed only in P276 (see **Figure 5H**), we focused our analysis on this patient cell line. Across genes within PP-rescued regions, we observed a modest upward shift in expression (**Figure 6E-6F**). Although the aggregate change did not reach statistical significance, the small number of expressed genes in this subset (n = 47) likely limits power. Focusing on genes meeting a lenient rescue criterion (significantly downregulated in P276 vs S1 and significantly upregulated in P276+PP vs P276 at FDR < 0.1), we identified several developmentally relevant genes, including the G protein–coupled receptor ADGRA3, the homeobox transcription factor MEIS2, and HDAC9, which is implicated in cardiac muscle development (example locus in **Figure 6G**). Gene Ontology analysis of PP-rescued genes showed enrichment for nervous system development terms (**Figure 6H**), reinforcing our earlier observations that SWING regions are preferentially enriched for developmental pathways, particularly neurodevelopmental programs. The overall magnitude of rescue was limited, which is expected given the low baseline expression of developmental programs in non-differentiating LCL model, limiting the ability for detection of robust transcriptional restoration.

In summary, to this point we demonstrate that in K562 cells, disruption of nuclear speckles repositions SWING regions toward the nuclear lamina, where genes become repressed, accompanied by gains in repressive histone modifications. This aberrant lamina association and gene repression are also observed in a patient-derived model with haploinsufficiency of SON. In contrast, loss of Lamin A/C or pharmacological speckle activation by PP alleviates lamina tethering and potentially favors gene activation. (**Figure 6I**).

### 8. Pyrvinium pamoate rescues early neurodevelopmental gene expression in iPSCs with acute SON depletion

As discussed above, the genes located within the SWING region are strongly developmental, however the K562 and LCL patient cell models used in the above analyses are not in a developmental context limiting the capture of relevant transcriptional changes. We therefore sought to investigate transcriptional consequences of speckle disruption during development and potential restoration with PP. To do so, we developed an induced pluripotent stem cell (iPSC) model bearing endogenous FKBP12^F36V^-tagged SON, enabling its rapid degradation. After 24h of dTAG treatment, SON was nearly undetectable, shown by immunofluorescence (**Figure 7A**; broader view in **Supplementary Figure 10A**) and by Western blot (**Figure 7B**). Notably, SON depletion caused a marked change in SRRM2-labeled speckle morphology where speckles became significantly smaller and rounder (**Figure 7A**; quantification in **Figure 7C**, comparing between DMSO and dTAG 500nM). Additionally, consistent with reports that SON is essential for cell and organismal viability^18^, high-dose dTAG treatment (500 nM for 24 h) caused extensive cell-cycle arrest (Mitotic cells circled in **Figure 7D**; broader view in **Supplementary Figure 10B**; quantification in **Figure 7E**).

**Figure 7.**
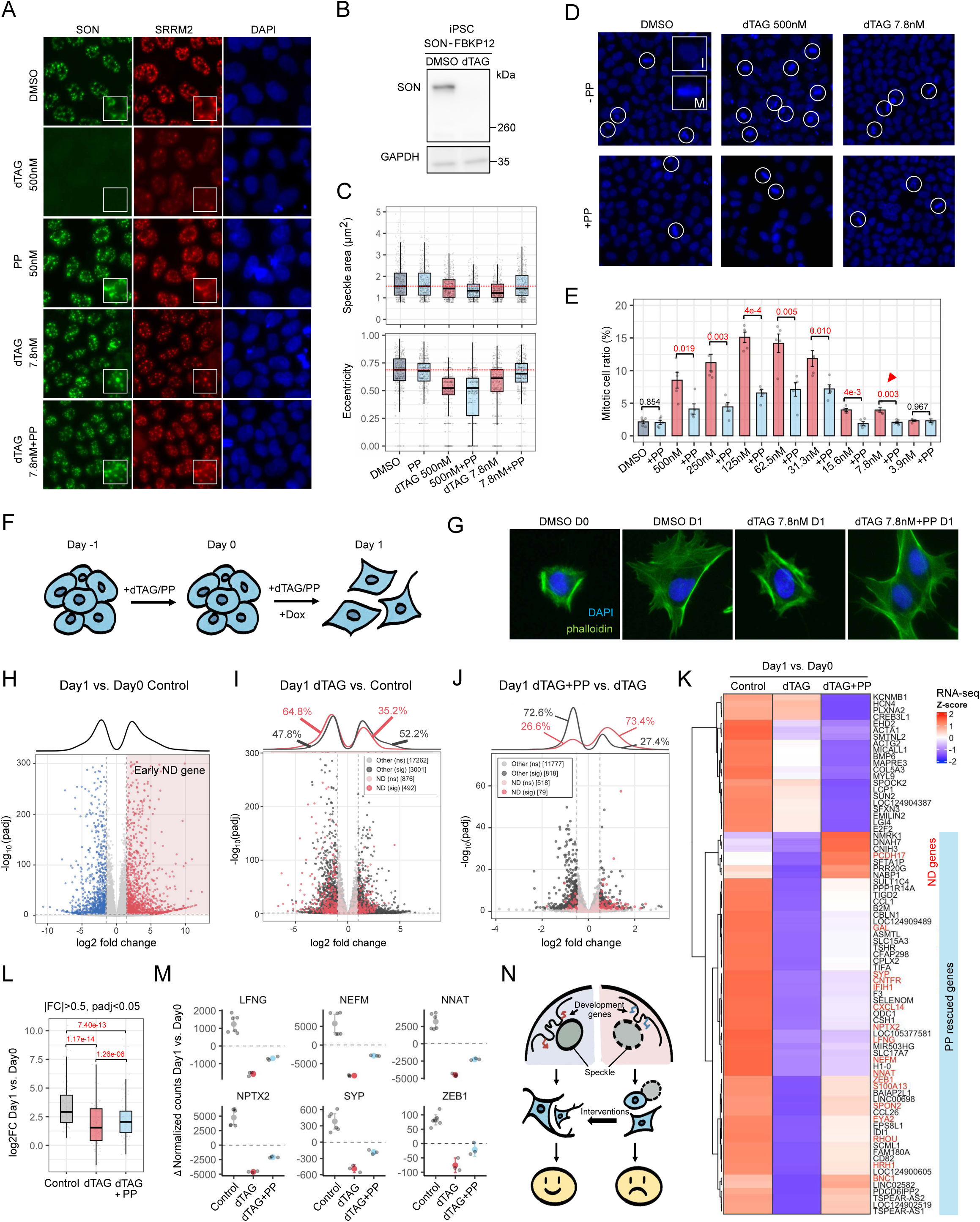
Acute SON depletion impairs transcriptional programs during iPSC to iNeuron differentiation and is rescued by pyrvinium pamoate. **(A)** Representative immunofluorescence (IF) images of iPSC SON–FKBP12 cells under the indicated treatments and staining conditions. Subpanel shows a zoomed-in view of a single cell. **(B)** Western blot for SON in iPSC SON–FKBP12 cells after 24 h treatments; GAPDH serves as a loading control. **(C)** Quantification of nuclear speckle area and eccentricity per nucleus from IF in (A). Box plots show median and interquartile range (IQR); whiskers, 1.5× IQR. P-values by two-sided Mann–Whitney U test. **(D)** DAPI images of iPSCs under the indicated treatments (PP = 50 nM). Metaphase cells are circled in white. **(E)** Fraction of metaphase cells under the indicated conditions. Error bars indicate SD. P-values by two-sided Mann–Whitney U test. **(F)** Schematic of the iPSC to early iNeuron differentiation timeline. **(G)** Representative immunofluorescence (IF) images of DAPI and phalloidin staining to visualize nuclei and actin filaments, respectively, showing cell morphology under the indicated conditions. **(H)** Volcano plot for RNA-seq: day 1 post doxycycline-induced differentiation vs. day 0. Highlighted genes meet FDR < 0.01 and |log2 fold change| > 1.5. Axes: x, log2 fold change; y, −log10(FDR). Top panel, density distributions for all differentially expressed genes. **(I)** Volcano plot for RNA-seq: day 1 dTAG vs. day 1 DMSO. Highlighted genes are early neuronal differentiation (ND) genes upregulated in (H) (log2 fold change > 1, FDR < 0.01). “Sig” denotes significantly differentially expressed genes (FDR < 0.05 and |log2 fold change| > 1). Top panel, density distributions for designated groups. **(J)** Volcano plot showing PP rescue in dTAG: rescue statistic R_PP = (dTAG+PP vs. dTAG) − (PP vs. DMSO). Highlighted genes are early ND genes whose induction is impaired by dTAG in (I) (log2 fold change < −1, FDR < 0.05). “Sig” denotes significantly rescued genes (FDR < 0.05 and |log2 fold change| > 0.5). Top panel, density distributions for designated groups. **(K)** RNA-seq heatmap of all significantly PP-rescued genes (FDR < 0.05 and |log2 fold change| > 1) with hierarchical clustering. Values plotted are the average Δ normalized counts under each treatment (day 1 − day 0). The y-axis lists individual genes. **(L)** Comparison of log2 fold change (day 1 vs. day 0) across conditions: control, dTAG, and dTAG+PP. for the PP-rescued gene set in (J). P-values by two-sided Wilcoxon signed-rank test. **(M)** Δ normalized RNA-seq counts (day 1 − day 0) for representative genes under the indicated conditions. **(N)** Model: speckle disruption impairs induction of developmental gene programs and neuronal differentiation, suggesting a potential therapeutic window for speckle-activating interventions (e.g., PP).

We next tested whether PP could mitigate the cellular consequences of SON depletion in iPSCs. PP alone did not affect speckle morphology or mitotic index (**Figure 7A, C–E**; DMSO vs. PP). Under conditions of near-complete SON depletion (500 nM dTAG), PP failed to restore speckle morphology **(Figure 7C, E**; 500 nM dTAG vs. 500 nM dTAG + PP), consistent with its proposed mechanism of modulating the biophysical properties of residual SON^39^.

In contrast, under partial depletion achieved by titrating dTAG, PP restored speckle morphology—both speckle area and shape (**Figure 7C; Supplementary Figure 10C–E**)—and reduced the mitotic-arrest phenotype in a dose-dependent manner (**Figure 7E**). Notably, at 7.8 nM dTAG, PP treatment restored both speckle morphology and mitotic progression to near-DMSO levels (**Figure 7D–E**).

We continued with partial depletion at 7.8 nM dTAG (as full depletion at 500 nM caused extensive cell cycle arrest) to examine whether SON depletion—and PP rescue—affects the developmental transcriptome. Because SON mutations cause ZTTK syndrome with a characteristic neurodevelopmental phenotype, we induced the iPSCs to neuronal differentiation using doxycycline activation of an NGN2 construct^42^ in SON–FKBP12 iPSCs. As outlined in the schematic (**Figure 7F**), iPSCs were pre-treated for 24 h with DMSO, dTAG, PP, or dTAG + PP on day -1 (D-1), followed by doxycycline induction of NGN2 on D0 to initiate neuronal differentiation; samples were collected at D0 and D1 for imaging and RNA-seq analysis. We did not extend the experiment beyond D2 because substantial cell death occurred even at 7.8 nM dTAG. On day 1 after NGN2 induction, control iPSCs exhibited morphological changes characterized by expansion of the cell body size and increased protrusion, consistent with a neural progenitor cells (NPC)-like character^42^ (**Figure 7G**, broader view in **Supplementary Figure 11A,** quantification in **Supplementary Figure 11B,** compare DMSO D0 and D1), while dTAG-treated iPSCs did not (**Figure 7G**, broader view in **Supplementary Figure 11A,** quantification in **Supplementary Figure 11B,** compare DMSO D1 with dTAG D1). Consistent with this, we observed robust induction of neuronal lineage and differentiation genes (hereafter referred to as early-ND genes), as well as inhibition of pluripotency genes in DMSO controls (D1 vs D0; transcriptome distributions in **Figure 7H**; example genes in **Supplementary Figure 11C**; Gene Ontology analysis in **Supplementary Figure 11D**).

We next investigated whether SON depletion affects expression of the early-ND genes. Comparing D1 dTAG with D1 DMSO cells, we found the majority of differentially expressed early-ND genes (64.7%) were downregulated, whereas genome-wide DEGs were roughly evenly split between up- and downregulation (**Figure 7I**; compare early-ND vs other genes), indicating that loss of speckle integrity selectively impairs neuronal differentiation gene expression.

We then asked whether PP rescues the defects brought by dTAG treatment. Morphologically, PP treatment rescued both the cell size and level of protrusion (**Figure 7G**, broader view in **Supplementary Figure 11A,** quantification in **Supplementary Figure 11B,** compare dTAG D1 with dTAG+PP D1). At the transcriptomic level, we asked whether PP restored the impaired induction of early-ND genes (early-ND genes with FC < 0 in **Figure 7I**) by comparing dTAG+PP D1 with dTAG D1. Because PP is known to have other physiological functions (including inhibition of mitochondrial respiration^43,44^), we controlled for these non-speckle mediated effects of PP by calculating a PP-rescue contrast in the dTAG background (see Methods for details). Using this control, we found that the majority of early-ND genes (73.4%) that were differentially expressed between Day 1 dTAG+PP vs dTAG showed restored expression (**Figure 7J**, “ND”; heatmap of significant early-ND genes in **Figure 7K**; summary quantification in **Figure 7L**), whereas non-ND genes (“other”) did not show this trend. Notably, multiple dTAG-impaired and PP-rescued genes are established neuronal lineage factors (highlighted in red in **Figure 7K**), including LFNG (early neuronal commitment), NNAT (neurite outgrowth and early neurodevelopment), and ZEB1 (neural progenitor marker) (**Figure 7M**).

Collectively, these results indicate that SON depletion is associated with altered expression of early neuronal differentiation programs, and that PP treatment can partially restore aspects of these pathways during early stages of iPSC-to-iNeuron differentiation (**Figure 7N**).

## Discussion

In this study, we identify a novel class of genomic regions—which we term SWING regions—that are sensitive to speckle dysfunction: either acute depletion in engineered cells or by clinical patient-derived mutations in speckle components relocates these domains towards the nuclear lamina. This repositioning is accompanied by increased H3K9me3 and preferential downregulation of genes residing in SWING regions. Notably, the SWING region genes are enriched for developmental pathways, with additional context-dependent enrichment for morphogenetic and tissue-organization genes. We further show that the nuclear lamina component Lamin A/C competes with speckles for SWING positioning and regulation: we find that Lamin A/C loss shifts SWING away from the lamina and induces the opposite transcriptional response, indicating inverse control of the SWING region by speckles compared to Lamin A/C. Finally, as proof of principle, pharmacological speckle activation with pyrvinium pamoate (PP) partially corrects both nuclear architecture—by reducing aberrant lamina association—and expression of a subset of developmentally relevant genes in patient cells and in iPSC model with acute SON depletion. These findings suggest the possibility of drug-mediated modulation of nuclear body–chromatin interactions to mitigate specific consequences of speckle-related dysfunction (**Figure 7N**).

Our study identifies nuclear speckles as determinants of specific genome features and, for the first time, shows that disrupting speckles directly alters chromatin organization by increasing nuclear lamina association (**Figure 1**). Although speckles are long correlated with active chromatin, the evidence is largely correlational, with direct perturbation tests previously limited. Prior work used siRNA against SRRM2 in a mouse cell line^12^, or a 6-hour acute co-degradation of SON and SRRM2 followed by 3D-genome readouts (HiChIP)^11^ ; in these prior studies there were minimal detection of global changes in contacts. In contrast, we reveal that the effects of speckle disruption are not genome-wide but rather concentrated to a defined subset of regions—the SWING regions. This domain specificity, and the partial or short-term nature of earlier perturbations, likely explains the limited changes detected previously.

Our results demonstrate competition for chromatin association of SWING regions between nuclear speckles and the nuclear lamina (**Figure 5**). Building on the inverse correlation between speckle and lamina association observed across cell lines^6,27^, we show that this competition specifically involves lamin A/C, a major mediator of chromatin–lamina interactions. Notably, lamin A/C can exist in a nucleoplasmic pool^45^, potentially regulated by phosphorylation^46^ which could be relevant to the relocation of SWING regions. We also note several similarities between SWING regions and facultative lamina-associated domains (fLADs), a subset of LADs exhibiting greater plasticity. Both regions show greater cell-type specificity and are more developmentally regulated than canonical LADs^47^, suggesting potential shared molecular underpinnings. Overall, the molecular mechanisms governing the interplay and competition between SWING regions and LADs remain to be determined. We establish that, in addition to regulating speckle-associated (SPAD) genes, nuclear speckles also regulate non-speckle-associated genes within SWING regions via a distinct mechanism. Prior work shows that speckles facilitate expression of speckle-proximal genes such as p53 targets^31^ and HIF-2α targets^32^, potentially by enhancing splicing efficiency^7^. Our results extend this model: speckles also compete with nuclear lamina mediated repression of non-speckle-associated SWING genes, limiting deposition of repressive marks such as H3K9me3 (**Figure 2**). We also find pathway differences between SWING regions and SPAD: SPAD genes are enriched for housekeeping and stress-response functions, whereas SWING regions appear to be enriched for developmental pathways (**Figure 3G–H**). We therefore hypothesize that, unlike SPAD genes that may require high basal and/or rapid expression, SWING genes could occupy an intermediate regulatory state that is particularly sensitive to speckle-mediated control, potentially enabling finer spatial and temporal regulation characteristic of developmental programs.

Multiple reports implicate mutations in nuclear speckle components, including SON and SRRM2 in neurodevelopmental disease^13,15,16^. In a clinical context using patient-derived cell lines, we observed that speckle dysfunction caused by naturally occurring mutations also led to aberrant lamina association, and correlated repression of SWING genes (**Figure 5-6**). We further found that treatment with pyrvinium pamoate (PP)—a small molecule drug reported to function by modifying biophysical properties of an intrinsically disordered region (IDR) of SON^39^ —rescued aberrant lamina association, and was accompanied by partial and context-dependent improvements in transcriptomic readouts in both a ZTTK patient-derived cell line and an iPSC model with acute SON depletion. While transcriptome-level rescue by PP was less uniform and likely reflects the compound’s multifaceted cellular activities and limited specificity for nuclear speckles, these data nonetheless support the concept that modulating speckle-associated biophysical properties can influence downstream chromatin organization and gene regulation, even as substantial work will be required before any application. Our results further highlight the potential of small-molecule approaches, as well as related strategies such as peptides or nanobodies, to partially restore nuclear body function by tuning the biophysical properties of their core components in disease contexts.

## Acknowledgments

We thank Dr. Brian Liau and Dr. Shelby Roseman for generously providing the SON/SRRM2 double-dTAG K562 cell line, and Dr. Andrew Belmont and Dr. Bas van Steensel for generously providing the LMNA and LBR knockout K562 cells, and Dr. Ian Krantz for providing all LCL lines used in this study.

## Data Availability

The genomic datasets generated during this have been deposited in the Gene Expression Omnibus under accession numbers GSE324462 and GSE324463.

## Material and Methods

This research complies with all relevant ethical regulations for studies involving cultured human cell lines by University of Pennsylvania.

## Experimental methods

### Cell culture

K562 derivatives—SON/SRRM2 FKBP12 double knock-in, LMNA knockout (LMNA−/−), and LBR knockout (LBR−/−)—were maintained in RPMI-1640 containing 2 mM L-glutamine (Invitrogen, 11875085), 10% fetal bovine serum (FBS), and 1% penicillin–streptomycin at 37 °C with 5% CO₂. Lymphoblastoid cell lines (LCLs) were cultured in RPMI-1640 supplemented with 10% FBS and 1% penicillin–streptomycin under identical conditions. Human iPSCs were maintained on Matrigel-coated plates (Corning, hESC-qualified, 354277) in Essential 8 Medium (Gibco/Thermo Fisher Scientific, A1517001) with daily medium changes at 37 °C, 5% CO₂. Cells were passaged at ∼70–80% confluency using ReLeSR (STEMCELL Technologies, 05872) for clumps or Accutase for single cells (Innovative Cell Technologies, AT104), with 10 µM Y-27632 ROCK inhibitor added after thawing or single-cell passage (Tocris, 1254).

For acute speckle disruption, cells were treated with 500 nM dTAGV-1 (Tocris 6914) for 24 h unless otherwise indicated; matched DMSO (Sigma D8418) vehicle controls were included. For pharmacologic speckle activation, pyrvinium pamoate (PP, Sigma P0027) was applied at 100 nM for 24 h unless noted.

All experiments used independently grown and treated biological replicates. Cell line identity and genotype were confirmed by Sanger sequencing and/or immunoblotting against the relevant tagged proteins. Cultures were routinely screened and found negative for mycoplasma contamination at least every six months.

### iPSC to iNeuron Differentiation

iPSCs were differentiated to iNeurons on poly-D-lysine/laminin–coated plates (PDL, Sigma P0899-10MG; laminin, Life Technologies 23017-015). DMSO, PP or dTAG were added at Day -1. From Day 0–Day1, cells were transitioned to a KOSR-based medium (KnockOut DMEM 10829-018, KSR 10828028, NEAA 10370088, GlutaMAX 35050061, β-ME 21985023) and induced with dual-SMAD inhibition plus Wnt blockade (LDN-193189, DNSK 1062368-24-4, 100 nM; SB431542, Tocris 1254, 10 µM; XAV939, Stemgent 04-00046, 2 µM) together with doxycycline to activate NGN2 (Sigma D9891-1G, 2 µg/mL).

### Western blotting

Samples were lysed with RIPA buffer (150mM NaCl, 1% IGEPAL, 0.5% sodium deoxycholate, 0.1% SDS and 50mM Tris pH7.4). Protein concentration was determined by bicinchoninic acid assay (BCA) protein assay (#23227, Life Technologies). Samples were separated using precast 4-12% Bis-Tris polyacrylamide gels (ThermoFisher Scientific, NP0321BOX) in the presence of SDS. Proteins were transferred to nitrocellulose membrane, blocked with 5% milk TBST, and probed overnight at 4 °C with designated primary antibodies (1:1000), including SON (Abcam, ab121759), SRRM2 (Sigma-Aldrich, S4045), Lamin B1 (Abcam, ab16048), Lamin A/C (Santa Cruz, sc-376248), histone H3 (Active Motif, 39763), GAPDH (Cell Signaling, 5174), and β-Actin (13E5; Cell Signaling, 4970L).Following TBST wash steps, membranes were incubated in secondary goat anti rabbit antibody (Goat Anti-Rabbit IgG (H + L)-HRP Conjugate, 1:10000, Bio-Rad, 1706515) for one hour, washed with TBST, and detected using using Thermo Scientific SuperSignal West Pico PLUS Chemiluminescent Substrate (ThermoFisher Scientific, 34577). Signal quantification was performed using ImageJ software.

### Immunofluorescence and antibodies

Cells were washed twice with PBS and fixed with Paraformaldehyde solution 4% in PBS (Santa Cruz Biotechnology, CAS 30525-89-4) for 10 minutes at room temperature, washed again with PBS (Corning, 21-031-CV) then permeated with 0.1% PBS Triton X-100 for 15 minutes. Cells were incubated overnight at 4 °C with primary antibodies diluted 1:200 in PBS, including SON (Abcam, ab121759), SRRM2 (Sigma-Aldrich, S4045), Lamin B1 (Abcam, ab16048), RBM25 (Sigma-Aldrich, HPA003025), SF3A66 (Abcam, ab77800), H3K9me3 (Active Motif, 39161), and H3K27me3 (Cell Signaling, 9733). Secondary antibody conjugated with Goat anti-Rabbit IgG Alexa Fluor™ 488 (ThermoFisher Scientific, A-11008) and Goat anti-Mouse IgG Alexa Fluor™ 647 (ThermoFisher Scientific, A-21235) were diluted 1:200 in PBS and used respectively. After 1 hour incubation at room temperature, samples were washed and incubated in 1:10000 DAPI (Invitrogen, D1306, 5 mg/mL) for 5min. Imaging was performed with Nikon Eclipse Ti2 widefield microscope. Cells were imaged at focal plane with a 40x objective, and a deep depletion CCD camera cooled to between -70 and -80C.

### Stochastic Optical Reconstruction Microscopy (STORM) imaging

Immunofluorescence was performed as described previously and in^48^, using a 1:50 dilution of anti-H3 primary antibody (Active Motif, 39763) and a 1:50 dilution of Alexa Fluor™ 647–conjugated secondary antibody (Thermo Fisher Scientific, A-21235) in PBS. STORM imaging was performed as described in the literature^21^. Briefly, images were acquired on an ONI Nanoimager (Oxford Nanoimaging) controlled by NimOS software (v1.19.4) using the 640-nm laser (exciting AF647). Cells were imaged in buffer containing 0.1 M cysteamine (MEA; 77 mg/mL stock; Sigma-Aldrich, 30070-10G prepared in 360 mM HCl; Fisher, A508-P500), 5% (w/v) glucose (Alfa Aesar, A16828), and a 1% GLOX solution. The GLOX solution was prepared by dissolving 14 mg glucose oxidase (Sigma-Aldrich, G2133) in 50 µL catalase (20 mg/mL; Roche, 106810) and bringing to 200 µL with 10 mM Tris (pH 8.0; Invitrogen, 15568025) containing 50 mM NaCl (Fisher, S217-500). Imaging buffer was replaced every 90 min. Acquisition settings were 15-ms exposure for 30,000 frames at 30 °C with constant high-intensity illumination at 647-nm excitation.

### Oligopaint DNA-FISH

Sample slides were fixed with 4% PFA in PBS for 10 minutes at room temperature, washed with PBS and permeabilized with 0.5% Triton in PBS for 15 minutes. Subsequently, DNA-FISH was performed as described^49^. Oligopaint DNA-FISH probes to target genes were designed across a 50kb region centered on the transcription start site as described^50^. Imaging was performed with Nikon Eclipse Ti2 widefield microscope. Cells were imaged at focal plane with a 40x objective.

### CUT&RUN sequencing

CUT&RUN was performed as described previously with minor modification^23^. For each condition, nuclei from ∼6×10^5 cells were isolated in nuclear isolation buffer (10 mM HEPES-KOH pH 7.9, 10 mM KCl, 0.1% NP-40, 0.5 mM spermidine, 1× Halt protease inhibitor cocktail) and bound to concanavalin A–coated magnetic beads (BioMag Plus) pre-equilibrated in binding buffer (20 mM HEPES-KOH pH 7.9, 10 mM KCl, 1 mM CaCl₂, 1 mM MnCl₂). After bead binding, samples were split for antibody incubations with Lamin B1 (Abcam, ab16048), SON (Abcam, ab121759), H3K9me3 (Active Motif, 39161), H3K27me3 (Cell Signaling, 9733), or normal IgG control (Millipore, 06-371); binding buffer was replaced with primary antibody diluted 1:100 in blocking buffer (20 mM HEPES-KOH pH 7.5, 150 mM NaCl, 0.1% BSA, 0.5 mM spermidine, 1× Halt protease inhibitor cocktail, 2 mM EDTA). IgG (Millipore 06-371) was included as a negative control. Bead–nuclei were rotated overnight at 4 °C, washed in washing buffer (20 mM HEPES-KOH pH 7.5, 150 mM NaCl, 0.1% BSA, 0.5 mM spermidine, 1× Halt protease inhibitor cocktail), and incubated with pA-MNase on ice. Digestion was initiated by addition of CaCl₂ and carried out for 30 min on a pre-chilled metal block on ice. Reactions were stopped by adding STOP buffer (200 mM NaCl, 20 mM EDTA, 4 mM EGTA, 50 µg/mL RNase A, 40 µg/mL glycogen) with gentle vortexing, followed by fragment release at 37 °C for 10 min. Supernatants were collected, treated with Proteinase K at 70 °C for 10 min, and DNA was purified by phenol:chloroform:isoamyl alcohol extraction. Libraries were prepared from released DNA using the NEBNext Ultra II DNA Library Prep Kit for Illumina, size profiles were assessed on an Agilent TapeStation, and concentrations quantified with the NEBNext Library Quant Kit (E7630, NEB). Libraries were sequenced on an Illumina NextSeq 1000 using NextSeq™ 1000/2000 P2 XLEAP-SBS™ Reagent Kit with 61bp per read.

### Polyadenylated RNA-seq

Cells were lysed in TRIzol (15596018, Thermo Fisher Scientific) and snap frozen. RNA was then isolated using chloroform extraction, followed by Zymo Direct-zol RNA Miniprep Kits (Zymo Research, R2050), including DNA digestion with DNase. Poly(A)+ RNA was further isolated using double selection with poly-dT beads (E7490, NEB). RNA-seq libraries were prepared using the NEBNext Ultra II Directional Library Prep Kit for Illumina (E7760, NEB). The size of the libraries was determined using a Bioanalyzer, and their concentration was determined using the NEBNext Lib Quant Kit (E7630, NEB). Libraries were sequenced on an Illumina NextSeq 1000 using NextSeq™ 1000/2000 P2 XLEAP-SBS™ Reagent Kit with 61bp per read.

## Data processing and analysis

### Statistics & Reproducibility

Details of the statistical tests are documented in the sections below for each analysis. For all replicate experiments in this study, cells were grown and treated separately as biological replicates. No statistical methods were used to pre-determine sample sizes, but our sample sizes are similar to those reported in previous publications ^6,31^. For all image-based statistics apart from STORM (see corresponding section for details), two-sided Wilcoxon test was performed, and no assumption of data distribution was made.

### Immunofluorescence (IF) data processing and analysis

All IF images were taken at the focal plane with the autofocus function of Nikon NIS software (5.30.04) on a Nikon Eclipse Ti2 widefield microscope, except for the STORM imaging in Figure 1B and 5B. Subsequent image processing was performed with CellProfiler^51^ (version 4.2.1, https://cellprofiler.org/) to identify nuclear speckle and measure various size, shape and intensity parameters. Pipeline and notes are available on Github: https://github.com/Chalietia/CellProfiler/.

### DNA-FISH processing and analysis

Images were preprocessed (flat-field correction and background subtraction) and analyzed in CellProfiler (version 4.2.1) with custom pipelines available at https://github.com/Chalietia/CellProfiler. Nuclei were segmented from DAPI using adaptive thresholding and size/shape filters to remove debris and touching clumps. DNA-FISH foci were detected per nucleus using a Laplacian-of-Gaussian spot detector with intensity and size thresholds chosen from control images; nuclei harboring >4 foci (indicative of over-segmentation or multiplets) were excluded a priori. Distances were measured in pixels and converted to physical units using the system pixel size (0.16 µm/pixel). Distance distributions were summarized as medians with interquartile ranges and compared across conditions using two-sided Mann–Whitney–Wilcoxon tests; where multiple comparisons were performed, p-values were adjusted by the Benjamini–Hochberg method.

### STORM imaging analysis

Localization data were drift-corrected in NimOS software (v1.19.4). Localizations at the nucleus were manually segmented from the field of view. The local compaction maps of the STORM image data were generated using Voronoi tessellation as previously described^52^, where each Voronoi cell is pseudo-colored based on its respective area. In addition to the density maps, O-SNAP (Objective Single-Molecule Nuclear Architecture Profiler; https://github.com/LakGroup/O-SNAP) was performed on the H3 localization data as described in the literature^21^. Briefly, the pipeline extracts 144 features from single molecule localization data. The features describe properties such as nucleus morphology as well as global and local organization of the H3 signal. O-SNAP then conducts parallel, orthogonal analyses to perform feature exploration. A volcano analysis identifies features that change most dramatically between experimental groups based on fold change and statistical significance using Welch’s t-test with the Benjamini-Hochberg method for multiple test correction. Here, O-SNAP highlights the modeled LAD thickness; the process for generating these modeled LAD segments is originally described in literature^22^.

### CUT&RUN data processing

Sequencing reads were aligned to the human reference genome GRCh38/hg38 using Bowtie2 v2.3.4.1 with the following parameters: --very-sensitive-local --no-mixed --no-discordant -I 10 -X 1000 -x hg38. Alignments were filtered with samtools v1.1 (view -q 5), sorted and indexed, and PCR duplicates were removed with Picard v2.26 (MarkDuplicates, REMOVE_DUPLICATES=true). Reads mapping to ENCODE hg38 blacklist regions were excluded from all downstream analyses (hg38 blacklist v2). Signal tracks (bigWig/bedGraph) were generated with deepTools (bamCompare) using -bs 5000 --smoothLength 15000 --effectiveGenomeSize 2913022398 --exactScaling --normalizeUsing RPKM --scaleFactorsMethod None -bl <hg38_blacklist.bed>, comparing each target to its matched IgG control.

### CUT&RUN/DamID differential analysis and SWING region definition

Differential analysis followed a published sliding-window framework^31^ (step-by-step: https://github.com/katealexander/TSAseq-Alexander2020/tree/master/genomicBins_DiffBind) with minor modifications. Briefly, PCR-deduplicated BAMs (targets with matched inputs/controls) were quantified over 50-kb windows tiled across hg38 at a 5-kb step (1/10 window) using DiffBind v3.12.0 (https://bioconductor.org/packages/release/bioc/html/DiffBind.html) with input subtraction enabled. Custom blacklist regions (generated as described above) and ENCODE hg38 blacklist were supplied to DiffBind via dba.blacklist(). Library-size normalization used DBA_NORM_RLE. Windows showing significant SON signal change (FDR < 0.01, Benjamini–Hochberg) were split by direction and merged into contiguous intervals with BEDTools (merge) to yield differential domains. For SWING region calls, differential regions exhibiting a significant increase in Lamin B1 signal (FDR < 0.01) in K562 dTAG-treated versus control, or in patient (P276) versus healthy donor (S1), were designated as SWING regions. Genes were assigned to domains by intersecting domain intervals with transcription start sites; genes whose TSS fell within a differential domain were extracted using a Python script adapted from getGenesWithin.py in the referenced workflow.

### LAD and SPAD calling

Broad domains were called with SICER2 (v1.1) ^53^ on hg38 using the following assay-matched controls: matched IgG for CUT&RUN, no-primary control for TSA-seq, and no-Dam (Dam-only) control for DamID: sicer -t <signal.bam> -c <control.bam> -s hg38 -fdr 0.1 -w 2000 -g 20000. For SPADs, <SIGNAL> was SON CUT&RUN or SON TSA-seq libraries; for LADs, <SIGNAL> was Lamin B1 CUT&RUN or Lamin B1 DamID, as indicated. Resulting BED files were sorted and filtered to remove ENCODE hg38 blacklist regions. Consensus domains for each condition were defined by strict replication: domains present in all biological replicates were intersected with bedtools (intersect, optionally followed by merge to consolidate adjacent fragments), and the shared overlap was taken as the final set for downstream analyses. When LAD/SPAD domains overlapped SWING regions, SWING regions were given precedence and corresponding overlapping region is removed by bedtools subtract. Overlapping bases were removed from the LAD/SPAD sets using bedtools subtract followed by sorting.

### Gene ontology (GO) analysis

Gene sets comprising (i) genes of which TSSs reside within SWING regions and/or (ii) genes with significant differential expression by RNA-seq (as described in Results) were analyzed with STRING (https://string-db.org/; organism: Homo sapiens) using default enrichment settings. GO Biological Process terms were tested with STRING’s built-in over-representation analysis and Benjamini–Hochberg multiple-testing correction; terms with FDR < 0.05 were considered significant. To reduce redundancy, significant terms were clustered by similarity at a threshold of 0.8, and representative terms were reported. Unless fewer were available, the first 10 significant (non-redundant) terms are shown. Input identifiers were supplied as official gene symbols, and background was set to the STRING genome-wide reference.

### RNA-seq processing and differential analysis

Reads were aligned to the human reference genome GRCh38/hg38 with STAR v2.7.1a (https://github.com/alexdobin/STAR) using:--outFilterType BySJout --outFilterMultimapNmax 20 --alignSJoverhangMin 8 --alignSJDBoverhangMin 1 --outFilterMismatchNmax 999 -- outFilterMismatchNoverReadLmax 0.04 --alignIntronMin 20 --alignIntronMax 1000000.Alignments were quantified to RefSeq gene models (GRCh38) using htseq-count (v0.11.1) in union mode (--type exon --idattr gene_id -s no). Gene-level normalization and differential expression were performed with DESeq2 (v1.40.2) using default settings (size-factor normalization; Wald test; Benjamini–Hochberg adjustment). Unless otherwise indicated, genes with FDR < 0.05 and absolute log2 fold change >1 were considered differentially expressed. Statistical differences between log2 fold changes of predefined gene groups were assessed using two-sided Mann–Whitney–Wilcoxon tests; where multiple comparisons were made, p-values were adjusted by the Benjamini–Hochberg method.

### Identifying PP-rescued gene in iPSC to early iNeuron differentiation

For rescue analyses (day 1 only), RNA-seq counts were modeled with DESeq2 using within-day contrasts to estimate a PP-specific rescue effect while controlling for PP’s baseline impact. We defined the rescue contrast as 𝑅_pp_ = (dTAG+PP vs dTAG) − (PP vs DMSO), which measures the incremental effect of PP in the dTAG background after subtracting PP’s standalone effect. Positive 𝑅_pp_values indicate PP-mediated restoration of expression beyond its baseline action, whereas negative values indicate the opposite. Significance was assessed by DESeq2’s Wald test with Benjamini–Hochberg adjustment, and interpretations in the text and figures refer to this day-1 contrast.

---

**Supplementary Figure 1.** : **(A)** Western blot quantification for Figure 1A. ACTB serves as a loading control. Box plots summarize band intensities (median, IQR; whiskers, 1.5× IQR). P-values by two-sided Student’s t-test assuming equal variances. **(B-C)** Representative images (B) and quantification (C) of IF-derived granularity and radial coefficient of variation for four nuclear speckle–localized factors—SON, SRRM2, RBM25, and SF3A66—in K562 cells with or without 24 h dTAG treatment. Violin plots show distributions with embedded box plots (median, IQR; whiskers, 1.5× IQR). P-values by two-sided Mann–Whitney U tests. **(D)** Western blots for H3 and SRRM2 in K562 cells with or without dTAG treatment; GAPDH serves as a loading control. **(E)** STORM images of K562 SON/SRRM2 double knock-in cells with (top) or without (bottom) 24 h dTAG treatment; the color scale encodes log10(Voronoi polygon area). **(F)** Output from O-SNAP analysis of STORM datasets comparing cells with or without dTAG treatment, showing all significant terms (padj < 0.05).

**Supplementary Figure 2.** (A) Representative genome browser tracks for three biological replicates of DMSO-treated K562 cells (left) and pairwise Pearson correlation among the three replicates (right). **(B)** Representative genome browser tracks of Lamin B1 DamID and Lamin B1 CUT&RUN in K562 cells (left) and Pearson correlation between DamID and CUT&RUN signals (right). **(C)** Representative IF images of Lamin B1 in K562 cells with or without 24 h dTAG treatment. **(D)** Western blot for Lamin B1 in the indicated samples; GAPDH serves as a loading control. Box plots summarize band intensities (median, IQR; whiskers, 1.5× IQR). P-values by two-sided Student’s t-test assuming equal variances. **(E)** Percentage distribution of base pairs across SPAD, SWING, LAD, and unassigned genomic regions identified from K562 Lamin B1 Cut&Run data.

**Supplementary Figure 3.** (A) 2D density plot showing Lamin B1 level (x-axis) versus SON–TSA-seq signal (y-axis) for LADs, SWING regions, and SPADs. 2D density plots display kernel density estimates. **(B–C)** Multiplex DNA-FISH (MERFISH) in IMR90 cells quantifying foci-to-speckle distance (B) and foci-to-lamina distance (C) for loci in LADs, SWING regions, and SPADs. **(D)** 2D density plot of MERFISH measurements for loci classified as LAD, SWING, or SPAD. **(E)** Distribution of lamin-associated domains (LADs), speckle-associated domains (SPADs), and intermediate regions (not classified as LAD or SPAD) in baseline K562 cells, shown genome-wide (“All”) and within SWING regions on each chromosome. **(F)** Example browser view of Lamin B1 DamID signal across four cell lines; bottom red track shows the per-bin standard deviation across cell lines. BED tracks indicate domain calls: LAD (red), SWING (purple), SPAD (blue). **(G)** Variability of Lamin B1 signal across four cell lines, summarized as standard deviation within LADs, SWING regions, and SPADs. **(H)** Proportions of low-, mid-, and high-variability regions (bottom, middle, and top tertiles by standard deviation) within LAD, SWING, and SPAD categories. **(I)** Distribution (% of bases) of A1, A2, B1, and B2 subcompartments genome-wide (All) and within LADs, SWING regions, and SPADs in K562 cells.

**Supplementary Figure 4.** (A) Volcano plot of RNA-seq differential expression in K562 (dTAG 24 h vs DMSO). Genes with FDR < 0.05 and |log2 fold change| > 1 are considered significant. Axes: x, log2 fold change; y, −log10 FDR. **(B)** Comparison of expression levels for genes located in LAD, SWING, and SPAD in DMSO-treated K562 cells. Box plots show median and IQR; whiskers, 1.5× IQR. **(C)** Gene Ontology Biological Process enrichment for all genes located in SWING regions identified in K562 cells; selected development-related terms are highlighted.

**Supplementary Figure 5.** (A) Western blot for Lamin A/C in K562 WT and LMNA knockout cell; GAPDH serves as a loading control. **(B)** Representative genome browser tracks comparing reference SON TSA-seq and SON CUT&RUN signal generated in this study. **(C)** Pearson correlation between SON TSA-seq and SON CUT&RUN signals.

**Supplementary Figure 6.** (A) STORM images of 24 hr PP treated K562 cells; the color scale encodes log10(Voronoi polygon area). **(B)** Output from O-SNAP analysis of STORM datasets comparing cells with or without PP treatment, showing all significant terms (padj < 0.05).

**Supplementary Figure 7.** (A) Sanger sequencing of the SON locus in a healthy control and patient 276. Red arrows indicate the heterozygous point mutations. **(B)** Human Phenotype Ontology (HPO) terms annotated for patient 276. **(C)** Sanger sequencing of the SRRM2 locus in a healthy control and patient 246. Red arrows indicate the heterozygous point mutations. **(D)** Human Phenotype Ontology (HPO) terms annotated for patient 246. **(E)** RNA-seq normalized counts for SON transcripts in healthy control and patient 276. Box plots show median and IQR; whiskers, 1.5× IQR. **(F)** RNA-seq normalized counts for SRRM2 transcripts in healthy control and patient 246. Box plots show median and IQR; whiskers, 1.5× IQR. **(G)** Representative IF against SRRM2 in LCLs from a healthy donor, patient 276 (P276) and patient 246 (P246) with or without 100 nM PP. **(H)** Quantification of speckle area per nucleus for IF in the indicated cell lines and conditions. Box plots as in (E); P values by two-sided Mann–Whitney U tests.

**Supplementary Figure 8.** (A) Representative genome browser view of Lamin B1 CUT&RUN signal in healthy donor (S1), patient 246 (P246), and P246 treated with 100 nM PP. Blue arrows mark regions with decreased lamina association; red arrows mark regions with increased lamina association in the indicated sample. Bottom tracks indicate domain calls: LAD (red), SWING (purple), SPAD (blue). A zoomed-out view of the light-blue highlighted region is shown at right. **(B)** Metagene profiles and heatmaps of Lamin B1 CUT&RUN signal across all called LADs, SWING regions, and SPADs, in S1, P246, and P246 with 100 nM PP. Metagene lines show mean normalized signal; heatmaps display z-scored signal per region, centered at region midpoints and ordered by mean signal. **(C)** Gene Ontology Biological Process enrichment for genes located in SWING region identified in P276.

**Supplementary Figure 9.** : **(A)** Volcano plot of RNA-seq differential analysis between patient 246 (P246) and healthy donor (S1). Points are colored by domain localization (SPAD, blue; SWING, purple; LAD, red). “Sig” denotes significantly differentially expressed genes (FDR < 0.05 and |log2 fold change| > 1); “NS” denotes not significant. Axes: x, log2 fold change; y, −log10 FDR. The panel above shows density distributions for the designated gene groups. **(B)** Comparison of significantly differentially expressed genes (FDR < 0.05 and |log2 fold change| > 1) between P246 and S1, grouped by localization (SPAD, SWING and LAD). Box plots show median and IQR; whiskers, 1.5× IQR. P values by two-sided Mann–Whitney U tests. **(C)** Percentile distributions showing the proportions of significantly down-regulated (blue) and up-regulated (red) genes within SPAD, SWING or LAD.

**Supplementary Figure 10.** (A) Zoomed-out immunofluorescence images corresponding to Figure 7A for DMSO and dTAG (500 nM). **(B)** Zoomed-out DAPI images corresponding to Figure 7D. **(C–E)** Quantification of nuclear speckle area (C), extent (D), and eccentricity (E) per nucleus from IF across the indicated extended dTAG concentrations. Box plots show median and interquartile range (IQR); whiskers, 1.5× IQR.

**Supplementary Figure 11.** (A) Zoomed-out immunofluorescence images corresponding to Figure 7G. **(B)** Quantification of cell body size (Left) and level of cell protrusion (Right) from IF across designated conditions. Box plots show median and interquartile range (IQR); whiskers, 1.5× IQR. **(C)** Normalized RNA-seq counts for the indicated pluripotency marker genes (upper) and neuronal marker genes (lower) in DMSO-treated samples at day 0 and day 1. **(D)** Gene Ontology Biological Process enrichment for genes significantly upregulated (FDR < 0.01 and |log2 fold change| > 1.5) between day 1 and day 0 post dox treatment in DMSO-treated iPSCs. Neurodevelopment-relevant pathways are highlighted.

