## Supplementary figure for "SWING regions prime chromatin for nuclear speckle–mediated gene regulation"

Supplementary Figure 1

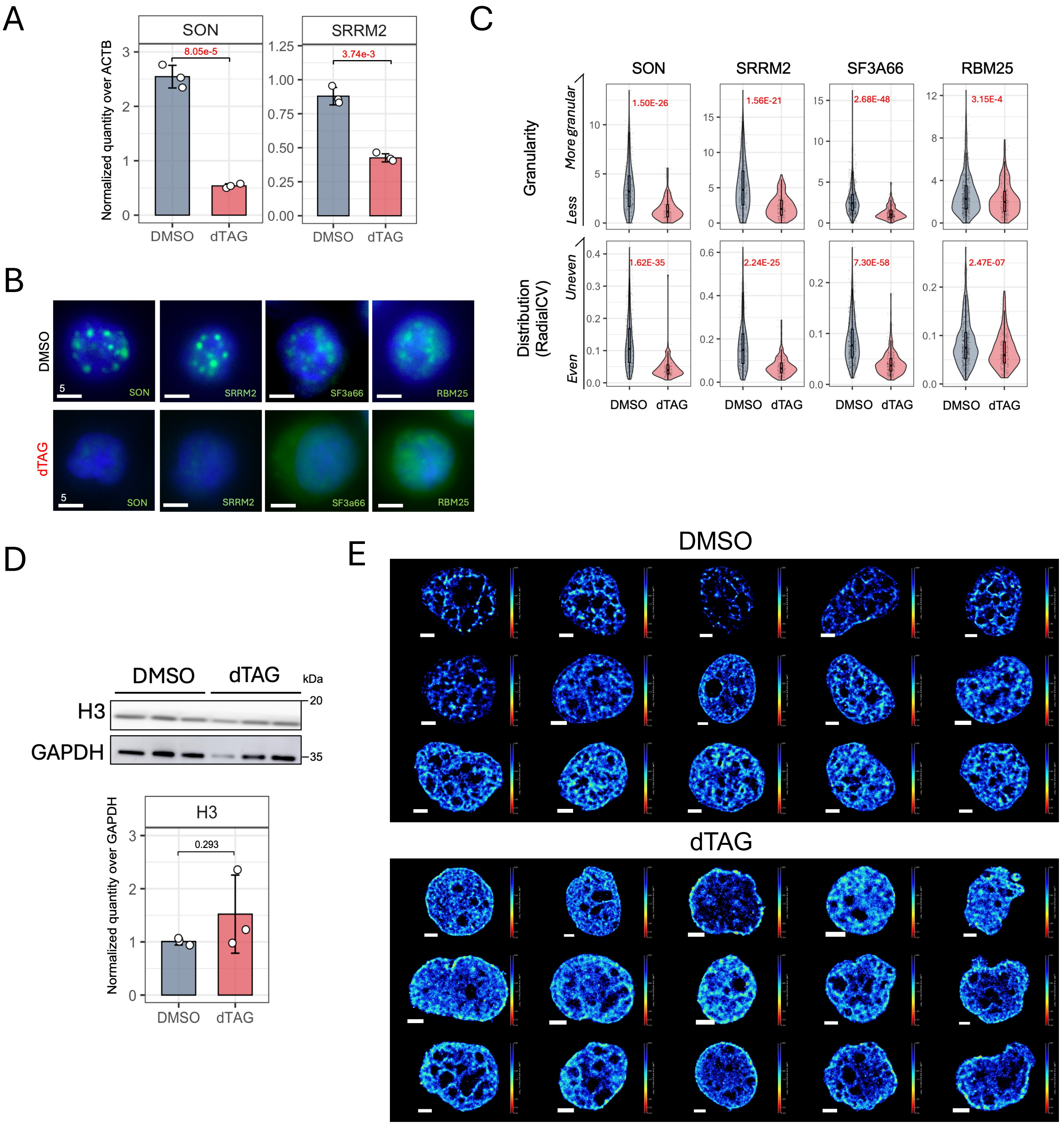


F

| Feature | DMSO | dTAG | Fold_change | log2_FC | adj_p_value |
|---|---|---|---|---|---|
| log_periphery_dbscan_cluster_radius_skewness | 0.369042 | 0.009705 | 0.026299 | -5.24886 | 0.018444 |
| log_interior_dbscan_cluster_spacing_skewness | -0.31398 | 0.014162 | 0.045103 | -4.47063 | 0.037323 |
| periphery_to_interior_dbscan_cluster_density_ratio | 1.008075 | 2.045623 | 2.029238 | 1.020938 | 5.96E-05 |
| **log_model_lad_segment_thickness_mean** | **0.465918** | **1.493279** | **3.205027** | **1.680337** | **2.67E-13** |
| model_lad_thickness | 21.55336 | 120.1768 | 5.575781 | 2.479174 | 1.22E-10 |
| periphery_dbscan_cluster_n_locs | 21808.16 | 143518.4 | 6.580947 | 2.718295 | 9.93E-09 |

Supplementary Figure 2

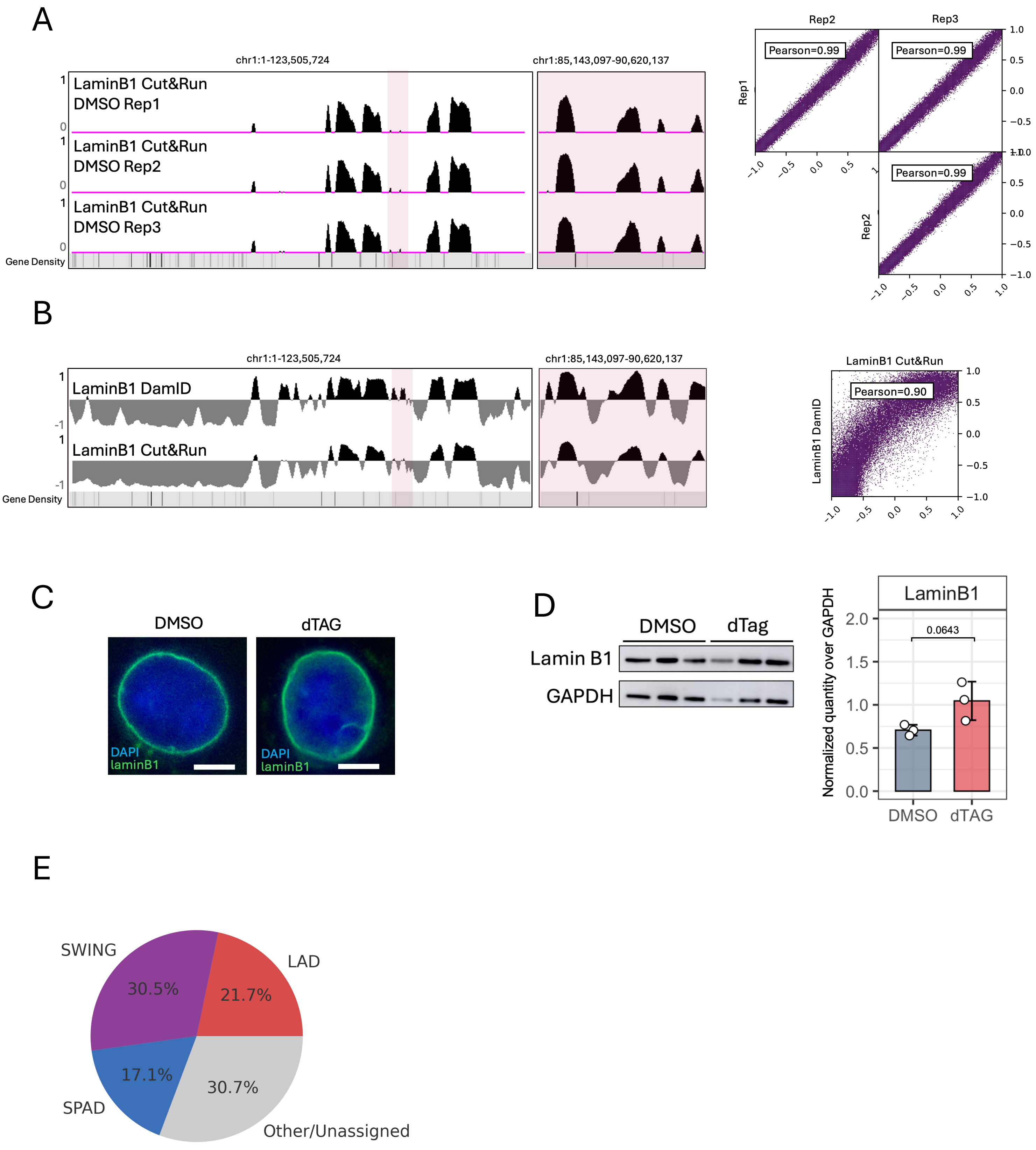

Supplementary Figure 3

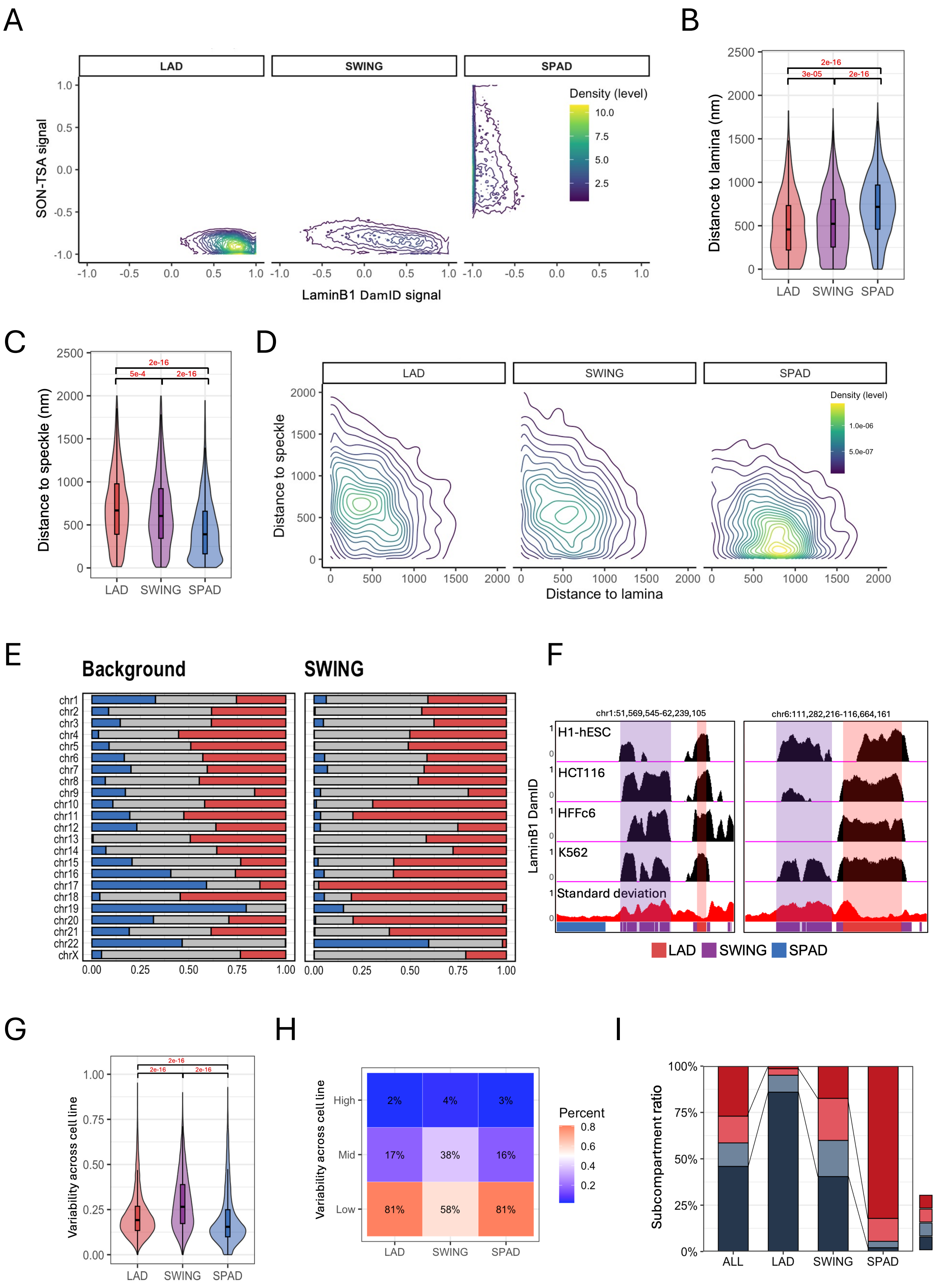

Supplementary Figure 4

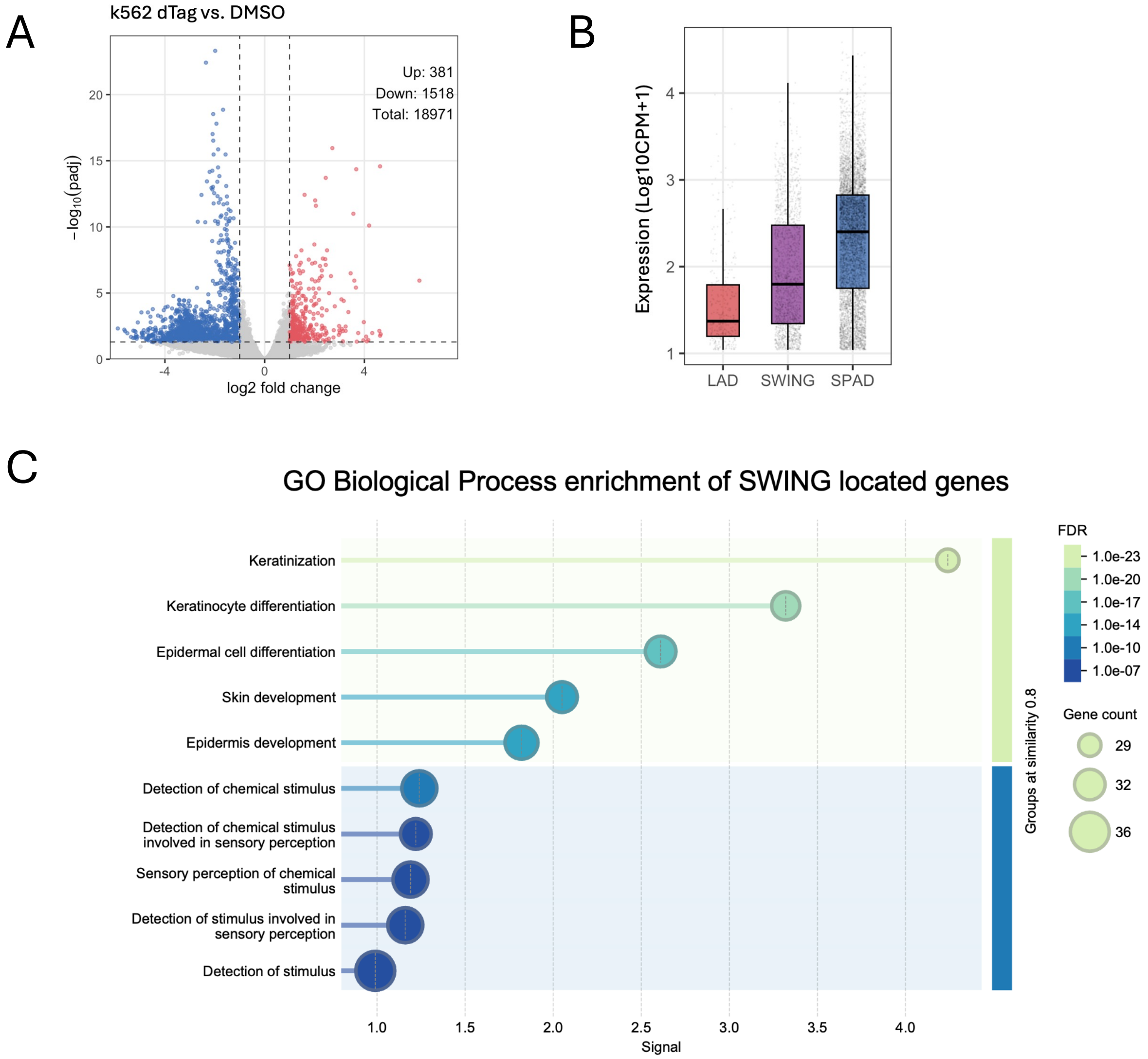

Supplementary Figure 5

A

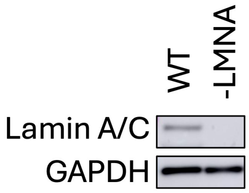


B

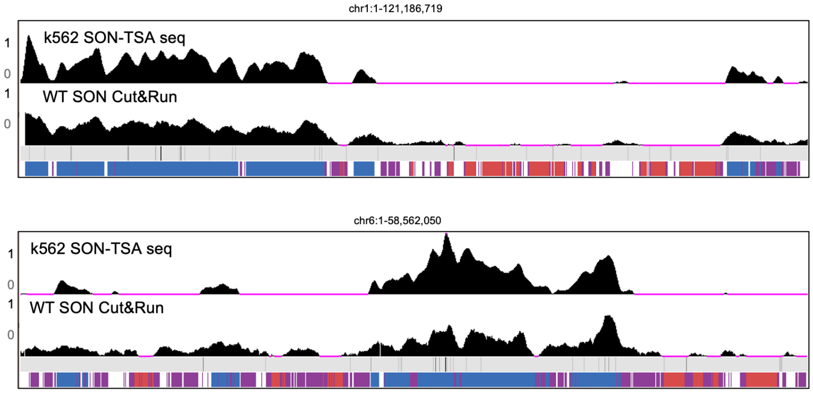


C

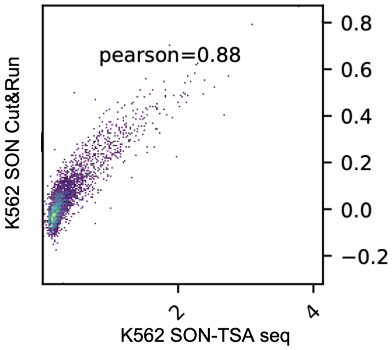

Supplementary Figure 6

A

100nM PP

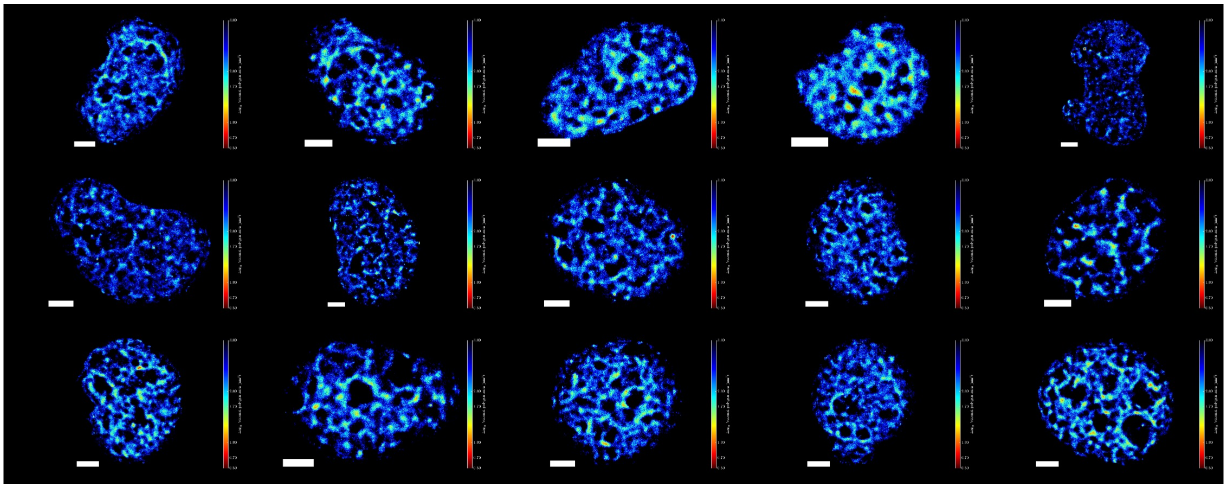

B

| Feature | DMSO | PP | Fold_change | log2_FC | adj_p_value |
|---|---|---|---|---|---|
| radial_loc_density_ring_gradient_major_axis | 1.67E-07 | 2.07E-08 | 0.124457 | -3.00628 | 0.016287 |
| radial_loc_density_ring_gradient_minor_axis | 2.08E-07 | 2.63E-08 | 0.126378 | -2.98418 | 0.016287 |
| radial_dbscan_cluster_density_ring_10 | 1.61E-06 | 6.73E-07 | 0.417975 | -1.25851 | 0.001924 |
| periphery_dbscan_cluster_n_clusters | 19.51613 | 8.157895 | 0.418008 | -1.2584 | 0.011542 |
| periphery_cluster_density | 2.05E-06 | 8.91E-07 | 0.434657 | -1.20205 | 0.000483 |
| **log_model_lad_segment_thickness_mean** | **0.465918** | **0.229879** | **0.493389** | **-1.0192** | **0.022738** |
| radial_dbscan_cluster_density_ring_09 | 2.22E-06 | 1.11E-06 | 0.499386 | -1.00177 | 0.000556 |
| log_model_lad_segment_thickness_skewness | 1.38708 | 2.867192 | 2.06707 | 1.047587 | 0.012367 |

Supplementary Figure 7

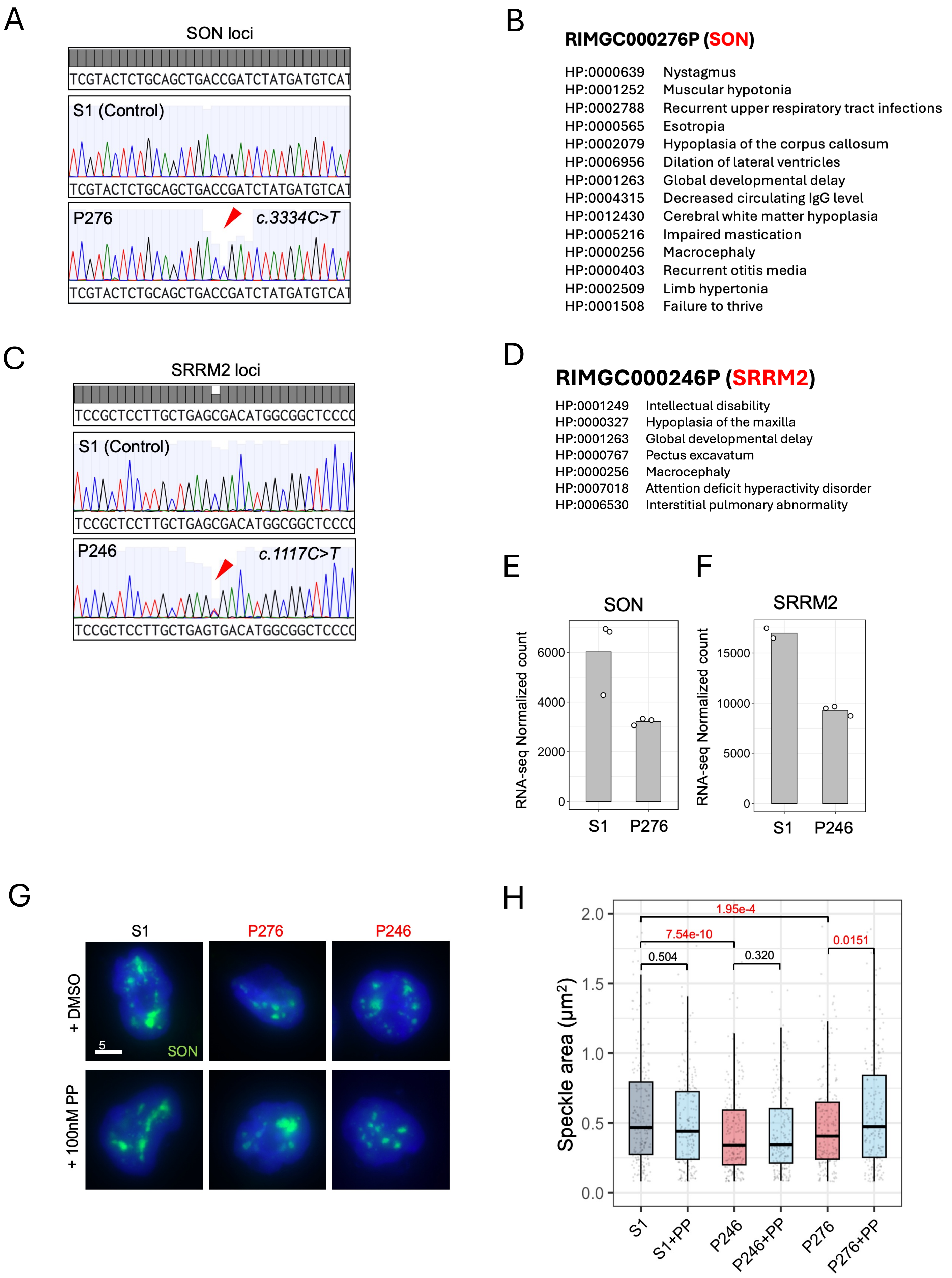

Supplementary Figure 8

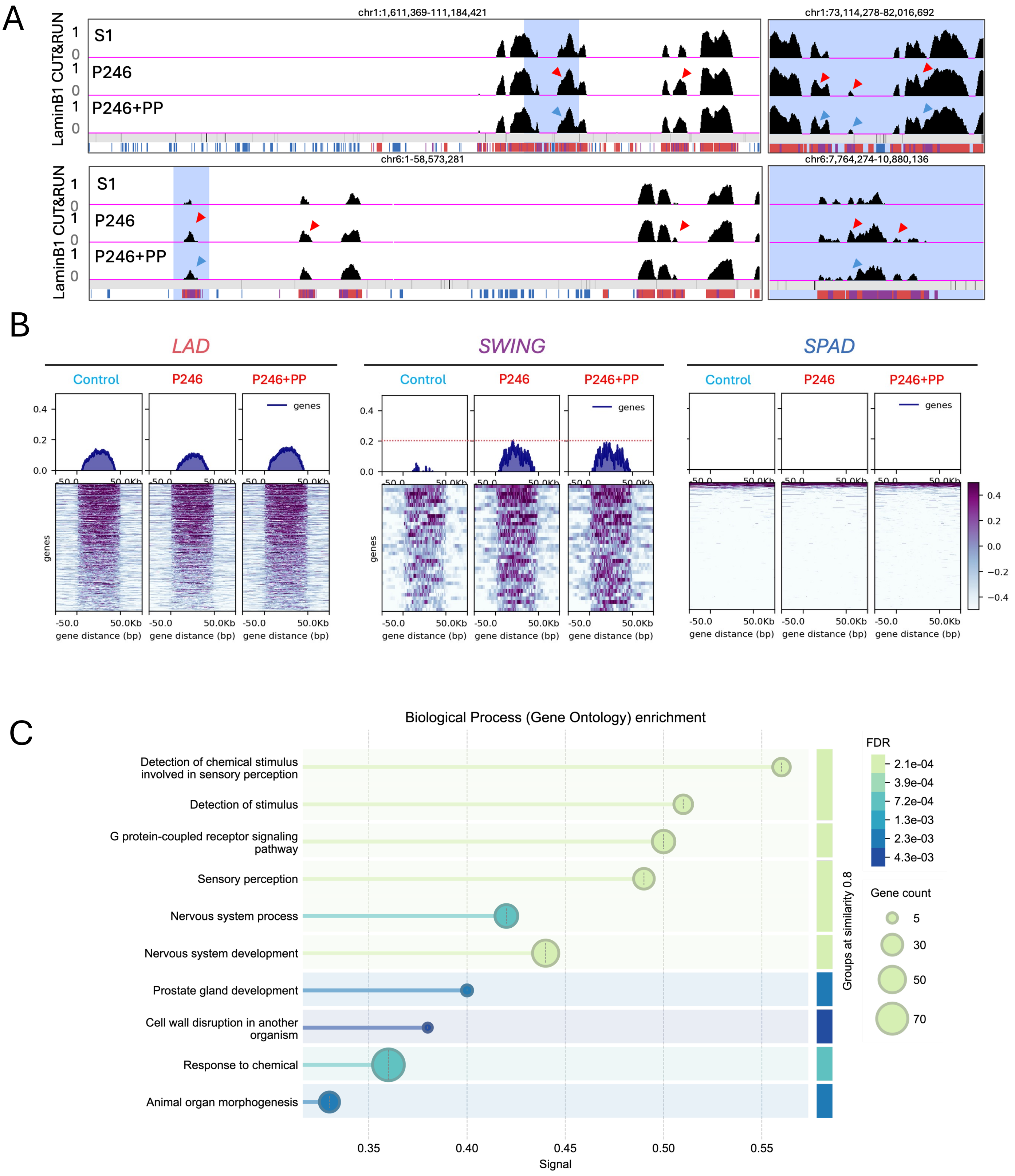

Supplementary Figure 9

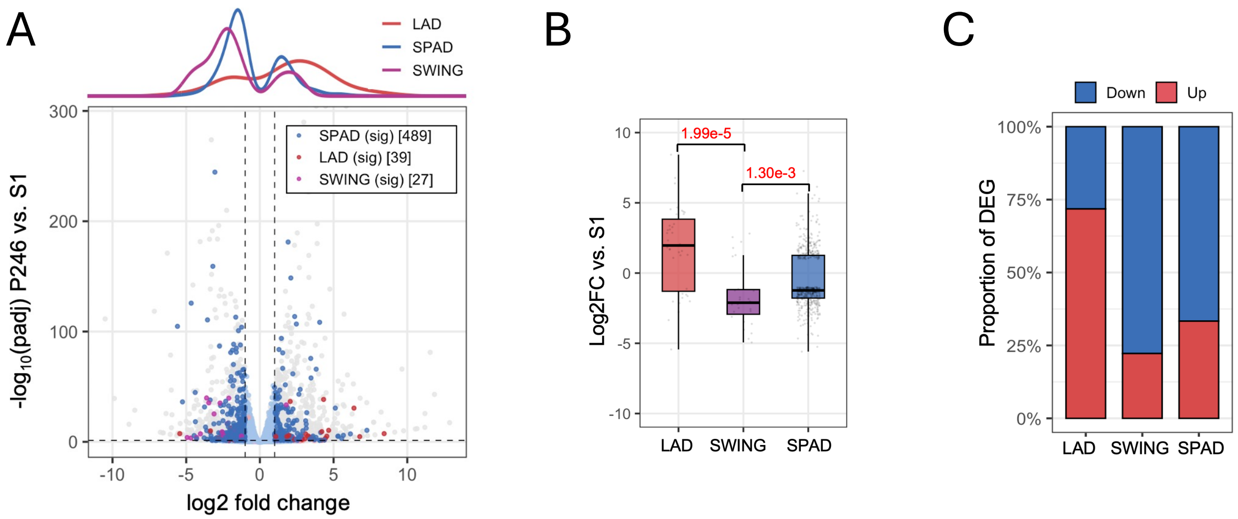

Supplementary Figure 10

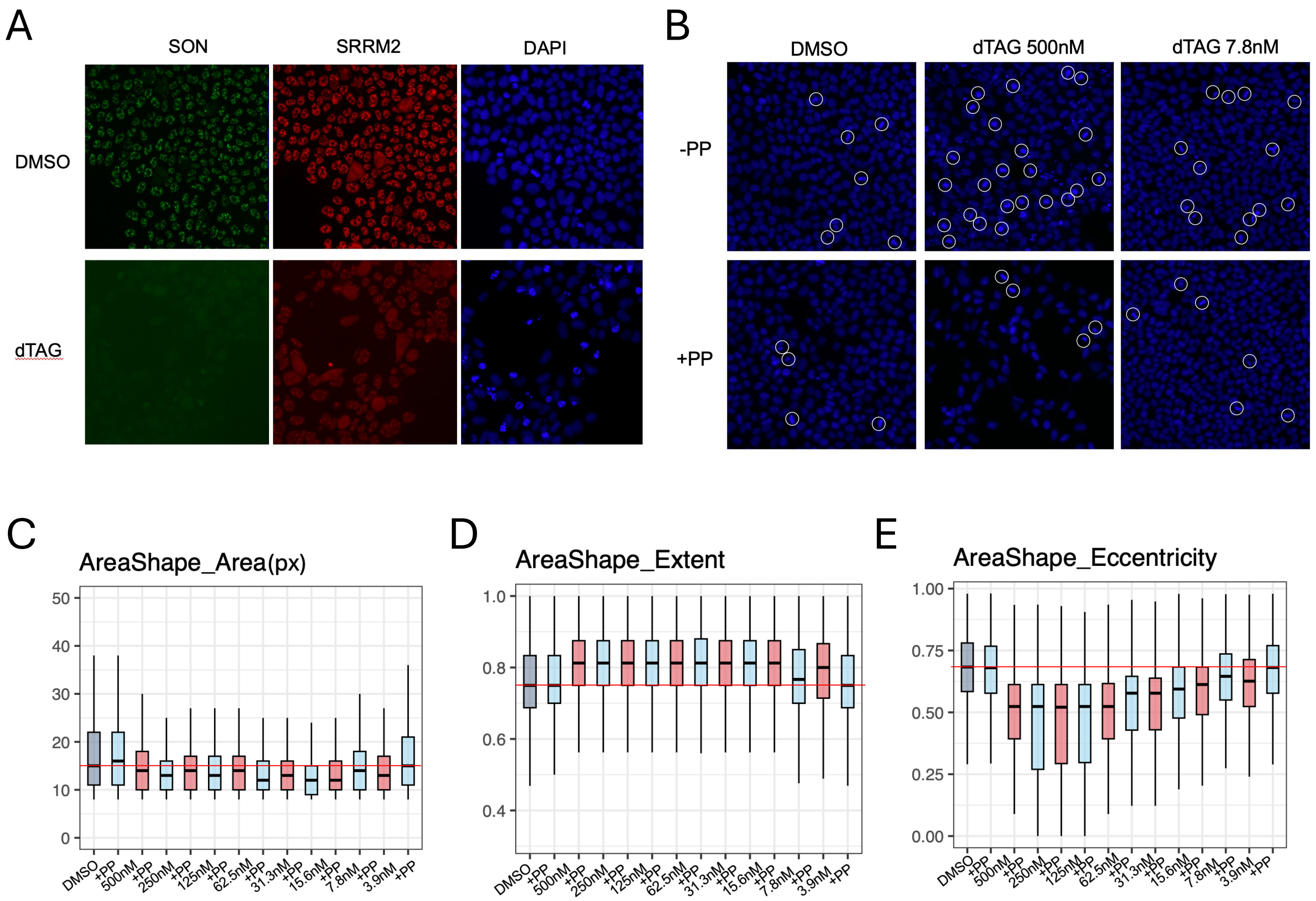

Supplementary Figure 11

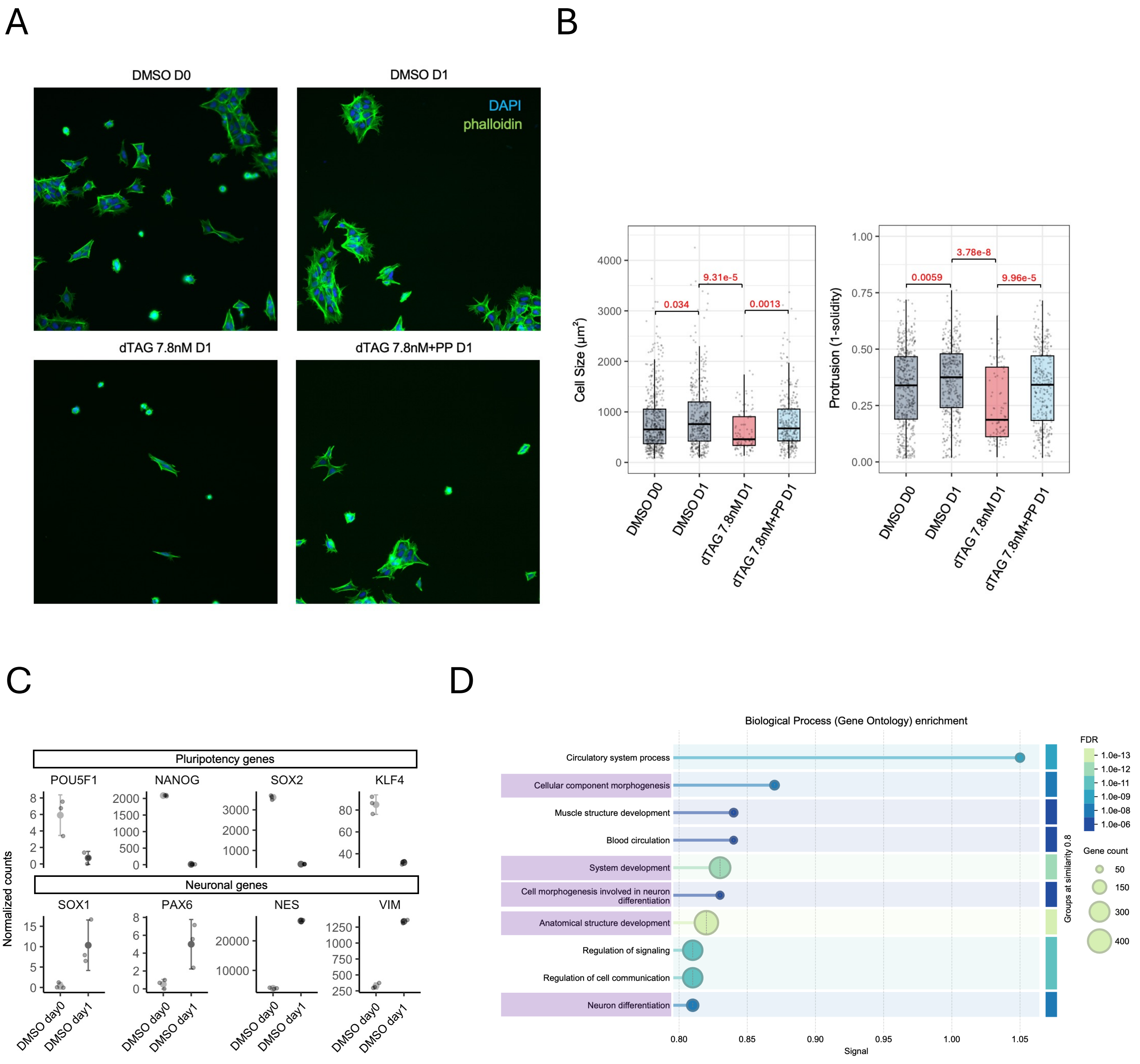
